# Ion channel inhibition by targeted recruitment of NEDD4-2 with divalent nanobodies

**DOI:** 10.1101/2024.05.28.596281

**Authors:** Travis J. Morgenstern, Arden Darko-Boateng, Emmanuel Afriyie, Sri Karthika Shanmugam, Xinle Zhou, Papiya Choudhury, Meera Desai, Robert S. Kass, Oliver B. Clarke, Henry M. Colecraft

**Author notes:** Department of Antibody Engineering, Genentech, Inc., South San Francisco, CA, USA. Equal contribution (Travis J. Morgenstern, Arden Darko-Boateng, Emmanuel Afriyie). Correspondence: Henry M. Colecraft, Department of Physiology and Cellular Biophysics Columbia University, College of Physicians and Surgeons 504 Russ Berrie Pavilion, 1150 St. Nicholas Avenue New York, NY 10032,; Oliver B. Clarke, Department of Anesthesiology, Columbia University, College of Physicians and Surgeons 1150 Saint Nicholas Avenue, New York, NY 10032.

## Abstract

Targeted recruitment of E3 ubiquitin ligases to degrade traditionally undruggable proteins is a disruptive paradigm for developing new therapeutics. Two salient limitations are that <2% of the ~600 E3 ligases in the human genome have been exploited to produce proteolysis targeting chimeras (PROTACs), and the efficacy of the approach has not been demonstrated for a vital class of complex multi-subunit membrane proteins— ion channels. NEDD4-1 and NEDD4-2 are physiological regulators of myriad ion channels, and belong to the 28-member HECT (homologous to E6AP C-terminus) family of E3 ligases with widespread roles in cell/developmental biology and diverse diseases including various cancers, immunological and neurological disorders, and chronic pain. The potential efficacy of HECT E3 ligases for targeted protein degradation is unexplored, constrained by a lack of appropriate binders, and uncertain due to their complex regulation by layered intra-molecular and posttranslational mechanisms. Here, we identified a nanobody that binds with high affinity and specificity to a unique site on the N-lobe of the NEDD4-2 HECT domain at a location physically separate from sites critical for catalysis— the E2 binding site, the catalytic cysteine, and the ubiquitin exosite— as revealed by a 3.1 Å cryo-electron microscopy reconstruction. Recruiting endogenous NEDD4-2 to diverse ion channel proteins (KCNQ1, ENaC, and Ca_V_2.2) using a divalent (DiVa) nanobody format strongly reduced their functional expression with minimal off-target effects as assessed by global proteomics, compared to simple NEDD4-2 overexpression. The results establish utility of a HECT E3 ligase for targeted protein downregulation, validate a class of complex multi-subunit membrane proteins as susceptible to this modality, and introduce endogenous E3 ligase recruitment with DiVa nanobodies as a general method to generate novel genetically-encoded ion channel inhibitors.

## Introduction

Induced proximity of enzymes to control the stability or function of target proteins is a rapidly evolving and powerful approach to not only develop new therapeutics for a broad range of diseases but also to devise enabling research tools (*1*). The premier example of this method recruits E3 ubiquitin ligases to foster targeted protein degradation (TPD) via the endogenous ubiquitin-proteasomal system (UPS) (*2, 3*). The most prevalent molecules for TPD fall into two classes — proteolysis targeting chimeras (PROTACs) are bivalent small molecules (or biologics, bioPROTACs) that simultaneously bind a target and an E3 ligase to form a ternary complex (*4*), and molecular glues, exemplified by the immunomodulatory drugs (IMiDs), thalidomide and derivatives, which were found by serendipity to act as degraders by binding and reshaping the surface of an E3 ligase to recognize a new target protein (*5–7*). TPD is either the underlying mechanism or has been successfully applied to conventionally undruggable targets including transcription factors IKZF1/IKZF3 (*5, 6*), androgen receptor (*8*), the anti-apoptotic protein BCL2L1 (*9*), and tau protein (*10*), with several molecules employing this modality currently in clinical trials, primarily for oncology indications (*11*). There are ~600 E3 ligases in the human genome, of which <2% have been exploited for TPD (*3*). The most advanced PROTACs and molecular glues exploit the E3 ligases von Hippel–Lindau tumor suppressor (VHL) and cereblon, respectively, which are both members of the cullin-RING family of E3 ligases (*3, 12*). The current reliance of TPD approaches on only a small number of E3 ligases is a recognized limitation as it may limit efficacy due to development of resistance or lack of expression of the specific E3 ligase in relevant tissues (*13, 14*). Therefore, expanding the universe of E3 ligases suitable for TPD is a critical endeavor in this nascent field (*3*).

To date, there has been scant application of TPD to cell surface ion channels, a multi-family class of membrane proteins that are necessary for diverse biological functions, including generating the heartbeat, neuronal firing, hormonal secretion, gastric acidification, fluid homeostasis, and sensory perception (*15, 16*). Ion channel blockers are essential therapeutics for a wide variety of diseases such as cardiac arrhythmias, pain, epilepsy, multiple sclerosis, and hypertension (*17*). Developing selective inhibitors for ion channels remains an incomplete and daunting task, typically approached by laborious high throughput screens for small molecule or toxin blockers of individual ion channels. TPD offers a potentially modular and general method to generate selective inhibitors for a broad spectrum of surface ion channels, but this approach is largely unexplored. Adaptations of the classical bioPROTACs design have been developed specifically for targeting integral membrane proteins (*2*). These include lysosomal targeting chimeras (LYTACs) (*18*), antibody-based PROTACs (AbTACs) (*19*), proteolysis targeting antibodies (PROTABs) (*20*), and transferrin receptor targeting chimeras (TransTACs). LyTACs and AbTACs employ antibody conjugates that simultaneously bind to an extracellular portion of the target protein and, respectively, to either the cation-independent mannose-6-phosphate receptor (CI-M6PR), a cell-surface to lysosome shuttling receptor, or to the rapidly recycling transferrin receptor. AbTACs and PROTABs also utilize extracellular bivalent antibodies that concurrently bind substrate protein and single-pass RING E3 ligases RNF43 and ZNRF3, respectively (*19, 20*). However, none of these approaches have been applied to ion channels, which typically are macromolecular complexes with multiple membrane-spanning segments and distinctive subunits that may make them challenging for TPD. Moreover, many ion channels have a large portion of their surface area in the cytosol with a comparatively small extracellular exposure, potentially making it challenging to develop extracellular antibodies for them (*21*).

Conceptually, it could be advantageous to exploit E3 ligases which are known physiological ion channel regulators for designing TPD approaches to this class of proteins. In this regard, the homologous to E6AP C-terminus (HECT) E3 ligase, NEDD4-2 (neural precursor cell expressed developmentally down-regulated 4-like) has been demonstrated to regulate the functional expression of numerous ion channels, including (but not limited to): voltage-gated Na^+^, Ca^2+^, and K^+^ channels; the epithelial sodium channel (ENaC); chloride channels (CFTR and ClC5); and water channels (AQP2) (*22*). Nevertheless, there are several obstacles to potentially exploiting NEDD4-2 for TPD of ion channels. NEDD4-2 (and NEDD4-1) are prototypes of the 9-member NEDD4 family of HECT E3 ligases which have a domain organization consisting sequentially of a C2 domain, 2-4 substrate-binding WW domains, and a catalytic HECT domain (*23*). To date, no member of the HECT E3 ligase family has been exploited for TPD, and their mechanism of action, which involves direct transfer of ubiquitin to a catalytic cysteine on the HECT domain before delivery to substrate, is qualitatively different from the cullin-RING E3 ligases. Further, there are no available suitable ligands of NEDD4-2 that can be leveraged for TPD. Finally, NEDD4-2 enzymatic activity is regulated by multi-layered mechanisms including phosphorylation (*24, 25*), protein-protein interactions (*26*), and auto-inhibitory intramolecular interactions (*27*), making it ambiguous whether its induced recruitment to ion channels would be effective in inhibiting their functional expression.

Here, we isolate a nanobody that binds selectively to a unique site on NEDD4-2 HECT domain in a manner revealed by cryo-electron microscopy (cryoEM) to not interfere with sites on the protein necessary for catalysis (the E2 binding site, catalytic cysteine, and the ubiquitin exosite) (*28*). Recruitment of endogenous NEDD4-2 to diverse ion channels — KCNQ1, ENaC, and Ca_V_2.2 — using a divalent (DiVa) nanobody format, in which the NEDD4-2 nanobody is coupled to a substrate targeting nanobody, resulted in a marked decrease in their functional expression to a similar extent as seen with simple NEDD4-2 over-expression. Global proteome analyses indicated DiVa-mediated recruitment of endogenous NEDD4-2 had minimal impact on cellular proteostasis, contrasting with changes in expression of hundreds of proteins observed with NEDD4-2 over-expression.

## Results

### Targeted recruitment of YFP-tagged NEDD4-2 HECT domain to reconstituted Ca_V_2.2 inhibits channel surface density and ionic currents

We used a heterologous expression system to test whether recruitment of NEDD4-2 HECT domain could down-regulate functional expression of a plasma membrane ion channel, Ca_V_2.2 (Fig. 1). The strategy was inspired by our previous finding that a construct consisting of a nanobody targeting the auxiliary β subunit of high-voltage activated calcium channels (nb.F3) fused to the NEDD4-2 HECT domain (Ca_V_-aβlator) potently inhibits Ca_V_1/Ca_V_2 channels (*29*). Ca_V_2.2 channels engineered with a double bungarotoxin binding site (BBS) in the extracellular domain IV S5-S6 loop (BBS-α_1B_) showed robust surface density by flow cytometry when co-expressed with Ca_V_β_2a_, YFP and individual nanobodies targeting these two proteins (nb.F3 and nb.YFP, respectively) (Fig. 1a, b; Supplemental Fig. 1). Accordingly, cells co-expressing α_1B_ + β_2a_ + YFP +nb.F3 + nb.YFP yielded robust whole-cell currents (Fig. 1c, d). Fusing nb.F3 and nb.YFP together with a flexible linker (GGGGSGGGGS) yielded a divalent (DiVa) nanobody, DiVa [nb.F3-nb.YFP], which had no impact on either reconstituted Ca_V_2.2 surface density (Fig. 1e, f; Supplemental Fig. 1) or whole-cell currents (Fig. 1g, h). Co-expressing reconstituted Ca_V_2.2, nb.F3 and nb.YFP with YFP-tagged NEDD4-2 HECT domain (YFP-HECT4-2) yielded modest decreases in Ca_V_2.2 surface density (Fig. 1i, j; Supplemental Fig. 1) and whole-cell currents (Fig. 1k, l; Supplemental Fig. 1). By contrast, co-expressing DiVa [nb.F3-nb.YFP] with reconstituted Ca_V_2.2 and YFP-HECT resulted in a significant decrease in channel surface density (Fig. 1m, n; Supplemental Fig. 1) and whole-cell currents (Fig. 1o, p; *I*_peak,0mV_ = −196.8 ± 44.8, *n* = 8 for α_1B_ + β_2a_ + α_2_δ-1 + nb.β + nb.YFP; and *I*_peak,0mV_ = −59.8 ± 18.5, *n* = 8 for α_1B_ + β_2a_ + α_2_δ-1 + DiVa [nb.F3-nb.YFP], *P* < 0.05, one-way ANOVA and Tukey’s multiple comparisons test). Overall, these results establish the proof-of-concept that recruiting NEDD4-2 HECT domain to an ion channel using DiVa nanobodies can effectively inhibit ionic currents by down-regulating channel surface density.

**Figure 1.**
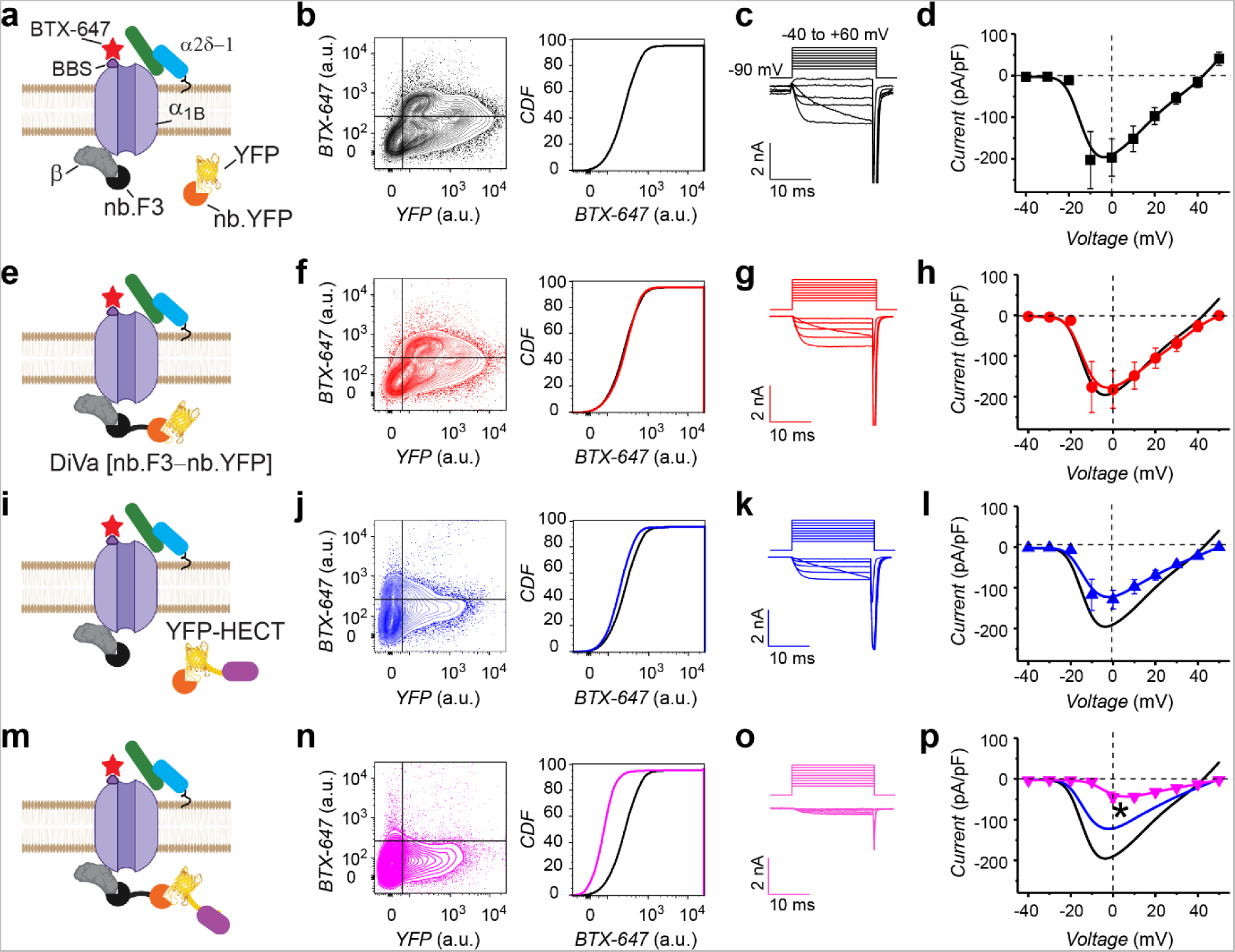
Targeted recruitment of YFP-tagged NEDD4-2 HECT domain to reconstituted Ca_V_2.2 inhibits channel surface density and ionic currents. (a) Schematic of BBS-α_1B_ + β + YFP + nb.F3 + nb.YFP. α_2_δ-1 was also co-transfected. Alexa Fluor 647-conjugated bungarotoxin (BTX-647) to an extracellular bungarotoxin binding site (BBS) engineered into α_1B_. (b) Exemplar flow cytometry contour and cumulative density frequency (CDF) plots. a.u., arbitrary units. (c) Exemplar whole-cell currents for cells expressing α_1B_ + β + α_2_δ-1 + YFP + nb.F3 + nb.YFP. (d) Population *I-V* curves. Data points are means ± SEM here and throughout; *n* = 8 for each point from three separate transfections. (e,f) Schematic and flow cytometry data for cells expressing BBS-α_1B_ + β + α_2_δ-1 + YFP + DiVa [nb.F3-nb.YFP]. (g,h) Exemplar currents and population I-V curves for cells expressing α_1B_ + β + α_2_δ-1 + YFP + DiVa [nb.F3-nb.YFP], *n* = 9 from three independent transfections. (i,j) Schematic and flow cytometry data for cells expressing BBS-α_1B_ + β + α_2_δ-1 + YFP-HECT4-2 + nb.F3 + nb.YFP. (k,l) Exemplar currents and population I-V curves for cells expressing α_1B_ + β + α_2_δ-1 + YFP-HECT4-2 + nb.F3 + nb.YFP. *n* = 8 from three independent transfections. (m,n) Schematic and flow cytometry data for cells expressing BBS-α_1B_ + β + α_2_δ-1 + YFP-HECT4-2 + DiVa [nb.F3-nb.YFP]. (o,p) Exemplar currents and population I-V curves for cells expressing α_1B_ + β + α_2_δ-1 + YFP-HECT4-2 + DiVa [nb.F3-nb.YFP], *n* = 9 from three independent transfections. \**p* < 0.05 by one-way ANOVA and Tukey’s multiple comparisons test.

### Identification and characterization of a nanobody that binds NEDD4-2 HECT domain

Given the lack of appropriate binders of NEDD4-2 that can be used for targeted recruitment of this E3 ligase, we sought to identify a nanobody that binds NEDD4-2 HECT domain (HECT4-2) but does not interfere with the catalytic activity of the enzyme. We expressed HECT4-2 bearing His- and FLAG-tags on the N- and C-terminus, respectively, in *E. coli*, and purified the protein from bacterial lysates using a nickel column (Fig. 2a, b). Purified HECT4-2 was used as bait to isolate nanobody binders from a yeast nanobody display library containing 1×10^8^ unique synthetic nanobodies presented on the extracellular surface of yeast (Fig. 2c; Supplemental Fig. 2) (*30*). Following two rounds of magnetic-activated cell sorting using 1 μM HECT4-2 as bait, individual clones were sorted into a 96-well plate, and their binding to HECT4-2 determined using flow cytometry (Supplemental Fig. 2). We obtained 15 unique nanobodies from our initial 96-well on-yeast staining assay, and herein focus on one positive clone, termed nb.C11. We cloned nb.C11 into a mammalian expression vector in frame with Cerulean (Cer) cDNA sequence. We co-expressed Cer-nb.C11 with Venus-tagged HECT4-2 (Ven-HECT4-2) in HEK293 cells and used flow cytometry fluorescence coupled with fluorescence resonance energy transfer (flow-FRET) (*31*) to probe binding between the two proteins in live cells (Fig. 2d). Cells expressing Cer.nb.C11 and Ven-HECT4-2 displayed robust FRET that was well fit by a binding curve, confirming interaction between the two proteins in live mammalian cells (Fig. 2d, g). By contrast, Cer-nb.C11 did not bind Ven-HECT4-1, despite the high homology between NEDD4-1 and NEDD4-2 HECT domains (83% sequence identity) (Fig. 2d, g).

**Figure 2.**
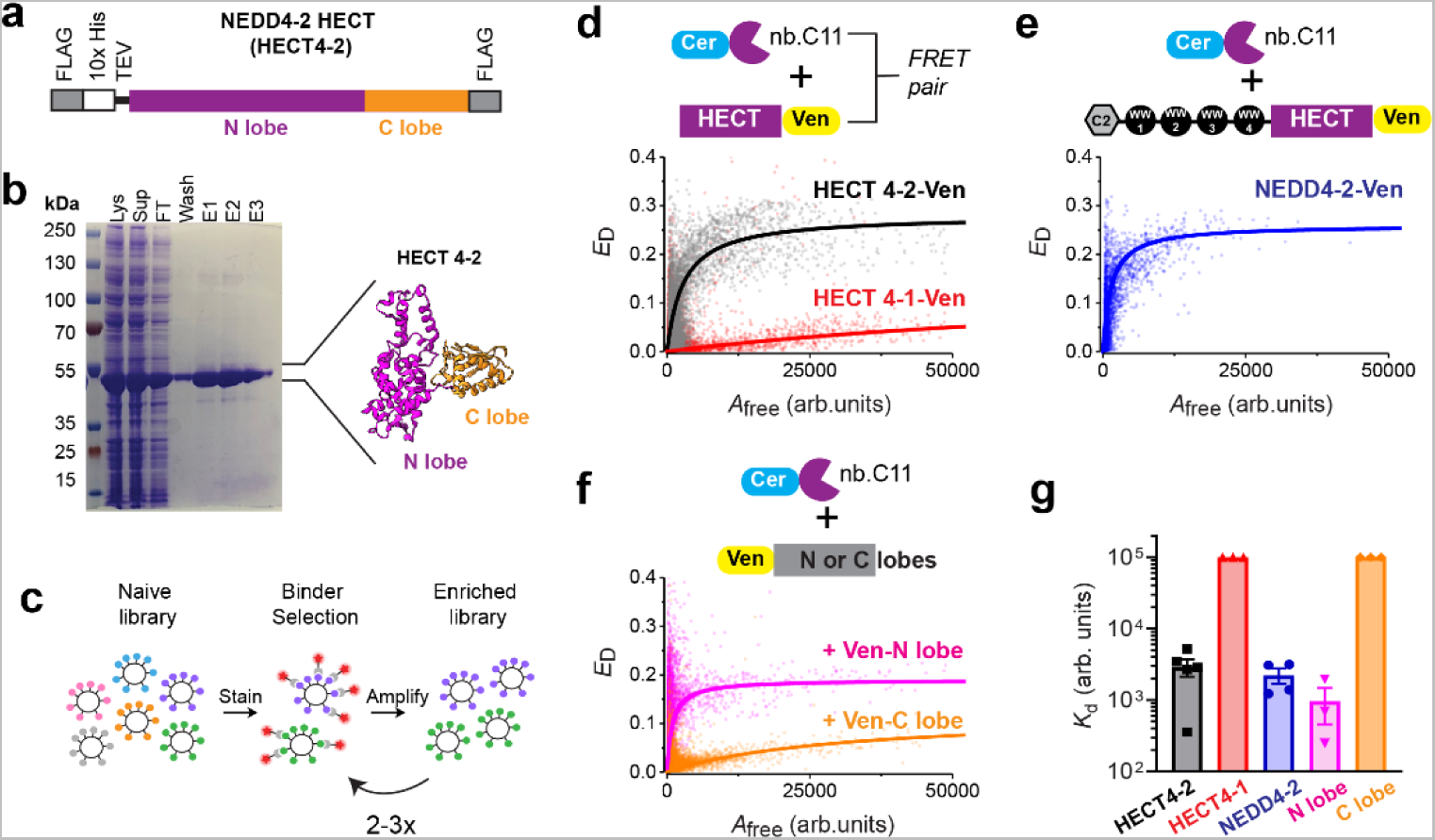
Identification and characterization of a nanobody that binds NEDD4-2 HECT domain. (a) Schematic of tagged NEDD4-2 HECT domain (HECT4-2) construct cloned into bacterial expression plasmid. (b) Coomassie-stained gel of purified HECT4-2. Inset: Structure of HECT4-2 domain (PDB ID: 3JW0) showing distinct N- and C-lobes in magenta and orange, respectively. Lys - cell lysate; Sup - supernatant; FT - flowthrough; E1-E3 - eluted fractions 1-3. (c) Schematic of yeast display selection method used to isolate nanobody binders to HECT4-2. (d) *Top*, FRET pair schematic, Cer-nb.C11 + Ven-HECT. *Bottom*, flow cytometry FRET (flow-FRET) scatter plots and binding curve fits for Ven-HECT4-2 and Ven-HECT4-1, respectively. (e) *Top*, FRET pair schematic, Cer-nb.C11 + Ven-NEDD4-2. *Bottom*, flow-FRET scatter plot and binding curve fit. (f) *Top*, FRET pair schematic, Cer-nb.C11 + either Ven-N-lobe or Ven-C-lobe. *Bottom*, flow-FRET scatter plot and binding curve fits for Ven-N-lobe and Ven-C-lobe, respectively. (g) Relative dissociation constants (K_d_) obtained from binding curve fits to flow-FRET scatter plots for binding interactions between nb.C11 and indicated FRET binding partners. a.u., arbitrary units.

Full-length NEDD4-2 contains a Ca^2+^-dependent membrane-targeting C2 domain and four substrate-binding WW domains upstream of the catalytic HECT domain. NEDD4 family E3 ligases are known to adopt intramolecular autoinhibitory interactions involving association of the HECT domain with WW and C2 domains (*32*). Thus, it was uncertain whether the epitope on HECT4-2 recognized by nb.C11 would be accessible in full-length NEDD4-2. Reassuringly, Cer-nb.C11 bound full-length Ven-NEDD4-2 with a similar affinity as the interaction with Ven-HECT4-2 (Fig. 2e, g). HECT4-2 has distinct N- (binds E2 enzymes and contains ubiquitin exosite) and C-lobes (contains the catalytic cysteine for ubiquitin transfer). Cer-nb.C11 preferentially bound HECT4-2 N-lobe (Fig. 2f, g). Thus, nb.C11 binds NEDD4-2, but not NEDD4-1, inside cells using an epitope on the HECT4-2 N-lobe that is accessible in the full-length NEDD4-2 protein.

### Structural basis of nb.C11 interaction with NEDD4-2 HECT domain

We turned to cryoEM to elucidate the structural basis for the selective interaction of nb-C11 with HECT4-2. Pilot experiments indicated that inclusion of the WW4 domain stabilized the NEDD4-2 HECT domain and provided additional ordered mass to facilitate structural studies (Supplemental Fig. 3). We assembled a complex of WW4-HECT4-2 and nb.C11 and confirmed nanobody binding by mass photometry and coelution on size exclusion chromatography (Supplemental Fig. 3). CryoEM analysis (Supplemental Figs. 4-7) resulted in a reconstruction with a global resolution of 3.1 Å (Fig. 3a), with minimal directional anisotropy (Supp. Fig. 4d). The average resolution in the nanobody was 3.1Å, and 3.16Å in the HECT-WW4 domain (Supp. Fig. 6). Consistent with flow-FRET data (Fig. 2f), nb.C11 was bound to the N-lobe of the NEDD4-2 HECT domain (Fig. 3). Density quality was sufficient to support generation and refinement of a near-complete atomic model, including confident assignment of the nanobody-HECT domain interface and the WW4 domain (Fig. 3). The WW4 domain forms a well-ordered fold with three anti-parallel β strands (β1-β3) (*33, 34*) interacting with a non-canonical PY motif (^589^PAVPY^593^) on the proximal loop connecting WW4 to the α1’ helix at the start of the HECT4-2 domain (Fig. 3; Supplemental Figs. 8,9). Thus, this ^589^PAVPY^593^ motif diverges from the canonical L/PPxY recognized by most WW domains of NEDD4-1 (*35, 36*) and NEDD4-2 (*33*).

**Figure 3.**
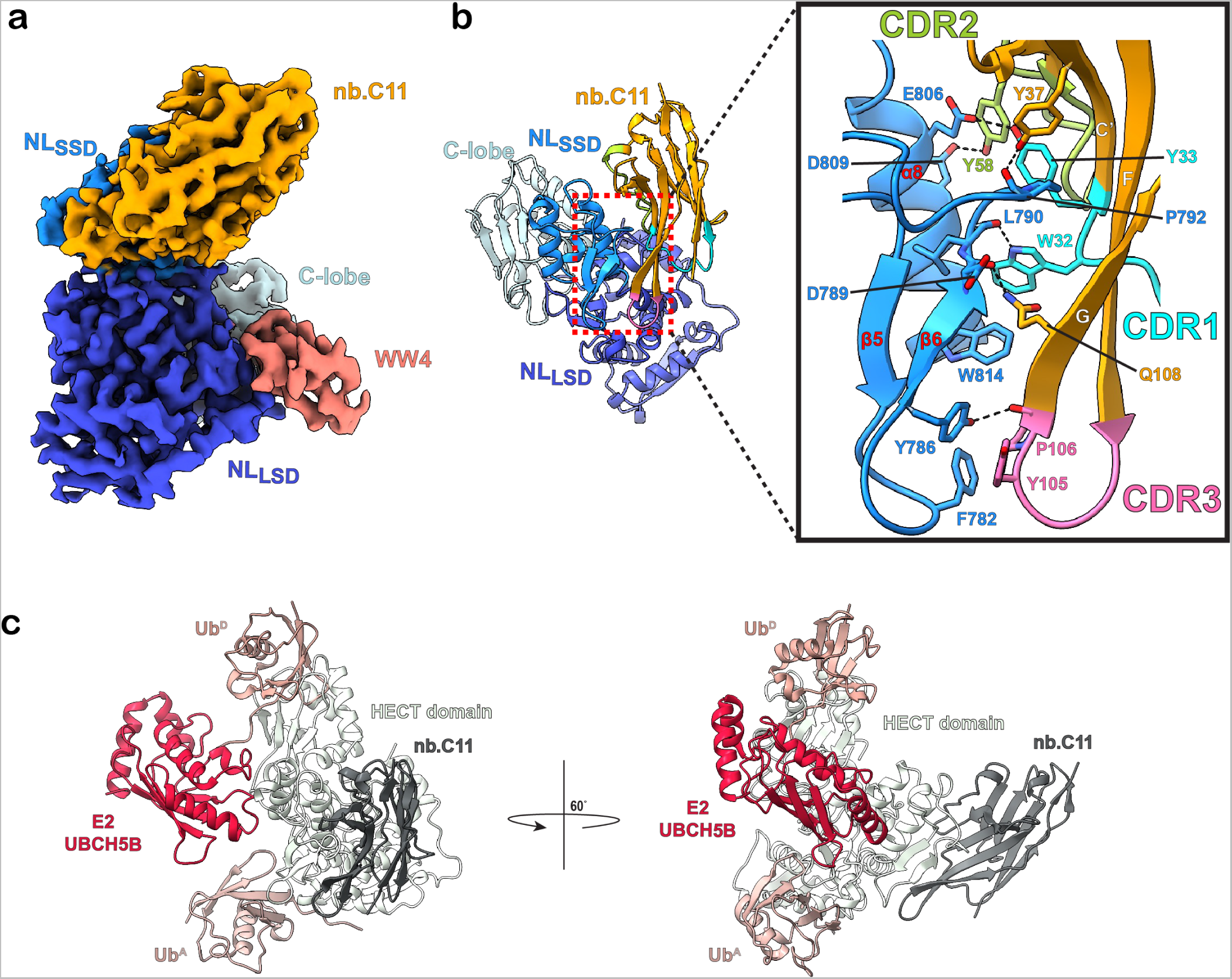
Cryo-EM structure of nb.C11 bound to HECT4-2 domain of NEDD4-2. (a) Final cryo-EM map of the WW4-HECT4-2:nb.C11 complex at 3.1 Å resolution. Sharpened density map is contoured at 0.124. (b) Atomic model of WW4-HECT4-2:nb.C11 complex including a close-up view showing NEDD4-2 N-lobe small subdomain (NL_SSD_) interactions with CDR1-3 loops on nb.C11. (c) Superposition of previously published E2 enzyme (UBCH5B) (PDB: 3JW0; red), donor ubiquitin (Ub^D^) (PDB: 3JW0; dark brown) and acceptor ubiquitin (Ub^A^) (PDB: 4BBN; light brown) binding sites on the HECT4-2:nb.C11 complex (HECT4-2, light gray; nb.C11, dark gray).

The HECT domain N-lobe is organized into two subdomains, referred to as the N-lobe large subdomain (NL_LSD_; includes α1’-α5 and α9-α10 helices; β1-β4 strands) and N-lobe small subdomain (NL_SSD_; comprises α6-α8 helices and β5-β6 strands), respectively (*37, 38*) (Fig. 3; Supplemental Fig. 9). Nb.C11 binds solely to the HECT4-2 NL_SSD_ (Fig. 3; Supplemental Fig. 9). Specifically, nb.C11 exclusively interacts with residues from the β5-β6 hairpin loop, the loop connecting β6 to α8, and α8 itself (Fig. 3; Supplemental Fig. 9). The binding interface is tightly packed and involves residues from all three nb.C11 complementarity-determining regions (CDR1-CDR3) and the scaffold (Fig. 3b). The interface buries surface area of 851.6 Å^2^ and contains a core region comprised of two asymmetric hydrophobic clamps. The first smaller clamp constitutes a hydrophobic boundary between HECT4-2 β5-β6 hairpin loop and nb.C11 CDR3, and involves face-to-face π-stacking between Y786 (HECT4-2) and Y105 (CDR3) and a CH/π face-to-face interaction (*39*) between Phe782 (HECT4-2) and P106 (CDR3). The Y786 sidechain also forms a hydrogen bond with the backbone carbonyl group of P106. The second larger clamp comprises W32 and Y33 (CDR1) forming π-stacking and hydrogen bonding interactions, respectively, with W814 and E806 from α8, while Y58 (CDR2) contacts D809 (α8) via a side-chain mediated hydrogen bond. W32 also forms a hydrogen bond with the backbone carbonyl of L790. In addition, the backbone carbonyl group from P792 (β6-α8 loop) forms a hydrogen bond and a CH/π interaction with Y37 (nb.C11 backbone). Finally, Q108 (nb.C11 backbone) forms a hydrogen bond with the D789 backbone carbonyl (Fig. 3b).

Crystal structures of E6AP and NEDD4-2 HECT domains in complex with UbcH7 and UbcH5B+ubiquitin, respectively, localized binding of the E2 conjugating enzymes to a hydrophobic groove on the NL_SSD_, primarily involving interactions with residues on the α7 helix (*38, 40*). The nb.C11 binding site is on the opposite face of the NEDD4-2 NL_SSD_ when compared to the E2 binding sites (Fig. 3c; Supplemental Fig. 9). This is a unique binding interface that has not been previously targeted by any ligand or observed as a site of protein-protein interactions, including with ubiquitin variants that bind NEDD4-2 HECT domain at either the E2 binding site or ubiquitin exosite (*28*). The nb.C11/WW4-HECT4-2 structure also provided insights into the basis for selectivity of nb.C11 for NEDD4-2 over NEDD4-1, despite the high sequence conservation in the NL_SSD_ between the two isoforms (Supplemental data Fig. 9). Of the eight residues on HECT4-2 involved in binding nb.C11, four are not conserved in NEDD4-1 (Y786H, D789E, P792N, and D809Y), and it is therefore reasonable to speculate that variation at these positions likely underlies the selectivity of nb.C11 for NEDD4-2 (Fig. 3b; Supplemental Fig. 9).

### Targeted recruitment of endogenous NEDD4-2 inhibits diverse channel types

KCNQ1 (Q1) pore-forming subunits tetramerize to form ion channels that are important for cardiac action potential repolarization and for salt and water transport in epithelial cells in organs including the lungs, cochlea, intestines, and kidneys (*41*). Q1 is known to be down-regulated by NEDD4-2 which binds a PY-motif located on Q1 C-terminus (*42*). Thus, we focused on heterologous expression of Q1 in HEK cells as a system to test the efficacy of DiVas featuring nb.C11 to recruit endogenous NEDD4-2. Control HEK293 cells transiently transfected with Q1-YFP and nb.YFP displayed large outward whole-cell currents in response to a family of test pulse potentials (from −60 mV to +100 mV in 20-mV increments) (Fig. 4, a-c). Co-expressing NEDD4-2 with Q1-YFP yielded substantially reduced whole-cell currents (Fig. 4, a-c), channel surface density (Supplemental Fig. 10, a-c), and Q1 protein expression (Fig. 4d), in accord with previous reports (*42*). Excitingly, we found that recruitment of endogenous NEDD4-2 to Q1-YFP using a divalent nanobody, DiVa [nb.YFP-nb.C11], yielded a similar suppression of whole-cell current amplitude (Fig. 4, a-c) and channel surface density (Supplemental Fig. 10, a-c) as was achieved with NEDD4-2 over-expression. Moreover, co-expressing DiVa [nb.YFP-nb.C11] also resulted in a reduced Q1-YFP expression assessed by Western blot (Fig. 4d), thereby correlating current suppression with cellular KCNQ1 levels.. Surprisingly, Q1-YFP currents reconstituted in Chinese hamster ovary (CHO) cells were unaffected by co-expressed DiVa [nb.YFP-nb.C11] (Supplemental Fig. 10g), in sharp contrast to the observations in HEK293 cells (Fig. 4c). We hypothesized that one potential explanation for this discrepancy was a difference in expression of endogenous NEDD4-2 between the two cell lines. Indeed, Western blotting showed that HEK293 cells express endogenous NEDD4-2 and the expression level is significantly up-regulated by transient transfection of exogenous NEDD4-2 (Supplemental Fig. 10h). By contrast, we could not detect expression of endogenous NEDD4-2 in CHO cells (Supplemental Fig. 10h,i). Our finding of a lack of NEDD4-2 expression in CHO cells is serendipitous, as it provides a critical control, showing that endogenous expression of NEDD4-2 is required for the functional effect of DiVa [nb.YFP-nb.C11] upon ion-channel functional expression to be evident.

**Figure 4.**
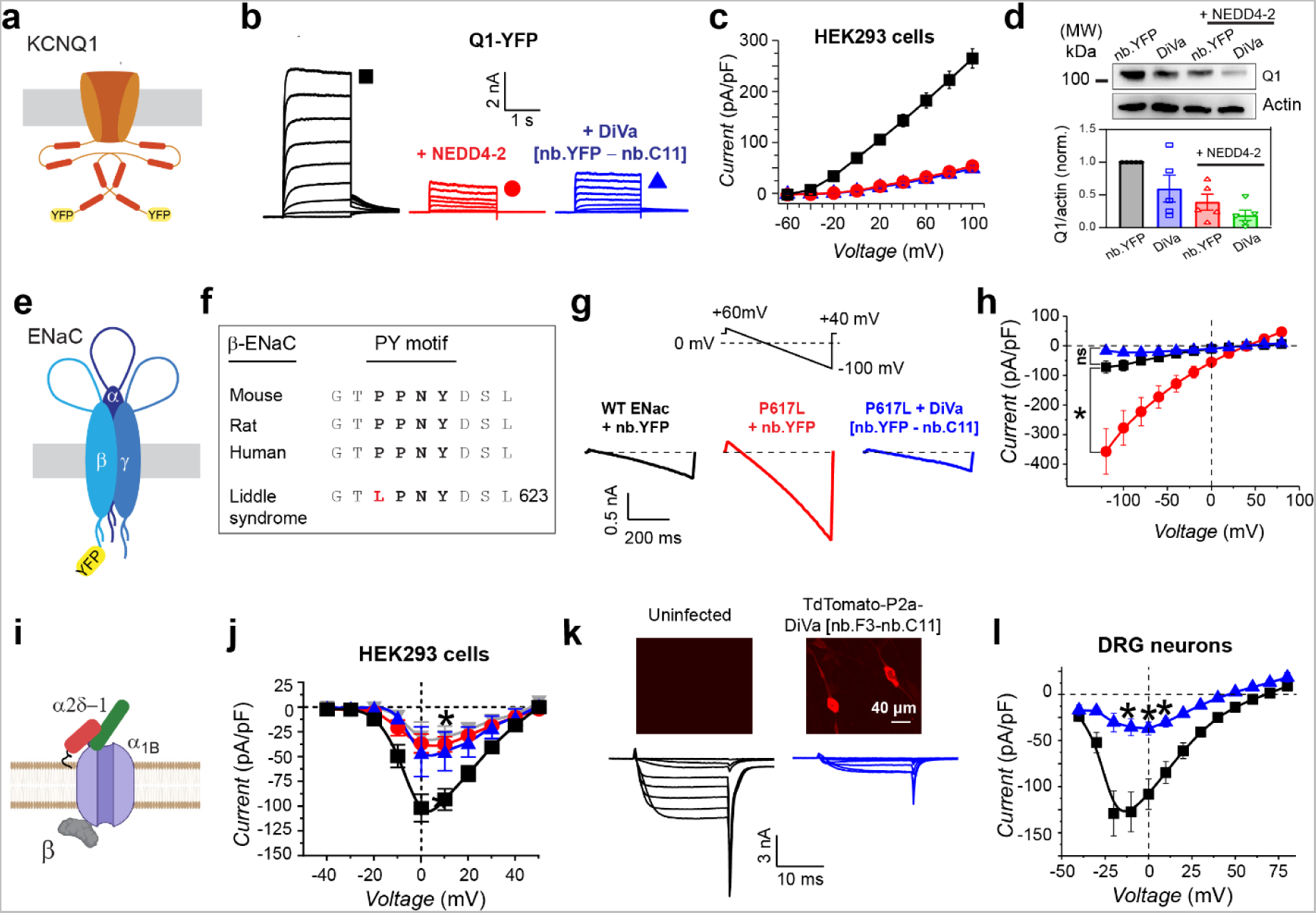
Targeted recruitment of endogenous NEDD4-2 inhibits diverse channel types. (a) Cartoon of KCNQ1-YFP (Q1-YFP). (b) Exemplar families of currents evoked from HEK293 cells co-transfected with Q1-YFP and nb.YFP (*left*); NEDD4-2 (*middle*) or DiVa [nb.YFP-nb.C11] (*right*). (c) Population *I-V* curves for Q1-YFP expressed with nb.YFP (black squares, *n* = 11), NEDD4-2 (red circles, *n* = 8), and or DiVa [nb.YFP-nb.C11] (blue triangles, *n* = 10). Data are means ± SEM from three independent transfections. (d) Exemplar Western blot (*top*) and population data (*bottom*) showing impact of NEDD4-2 and DiVa [nb.YFP-nb.C11] co-expression on Q1-YFP steady-state expression levels. (e) Cartoon of the epithelial sodium channel (ENaC) assembled with α, YFP-tagged β, and γ subunits. (f) Sequence alignment of conserved β-ENaC PY motif and showing position of a Liddle syndrome mutation, P617L. (g) *Top*, voltage ramp protocol. *Bottom*, exemplar ENaC currents from cells expressing WT ENaC + nb.YFP (*left*), ENaC[P617L] + nb.YFP (*middle*), or ENaC[P617L] + DiVa [nb.YFP-nb.C11] (*right*). (h) Population ENaC *I-V* curves evoked from HEK293 cells expressing WT ENaC + nb.YFP (black squares, *n* = 20), ENaC[P617L] + nb.YFP (red circles, *n* = 15), or ENaC[P617L] + DiVa [nb.YFP-nb.C11] (blue triangles, *n* = 9). (i) Cartoon of Ca_V_2.2 channel complex comprising α_1B_ + β_2a_ + α_2_δ-1 subunits. (j) Population *I-V* curves evoked from HEK293 cells expressing Ca_V_2.2 and nb.F3 (black squares, *n* = 32), NEDD4-2 (red circles, *n* = 28) or DiVa [nb.C11-nb.F3] (blue triangles, *n* = 9). Data are means ± SEM from at least three independent transfections. \**p* < 0.05 compared to nb.F3 by one-way ANOVA and Dunnett’s multiple comparisons test. (k) *Top*, confocal images of DRG neurons either uninfected (*left*) or infected with adenovirus expressing tdTomato-P2a-DiVa [nbF3-nbC11] (*right*). *Bottom*, exemplar families of whole-cell Ba^2+^ currents evoked by step depolarization protocols applied to DRG neurons infected with adenoviruses expressing mCherry (*left*) or tdTomato-P2a-DiVa [nb.F3-nb.C11] (*right*). (l) Population *I-V* curves from DRG neurons expressing mCherry (black squares, *n* = 8) or tdTomato-P2a-DiVa [nb.F3-nb.C11] (blue triangles, *n* = 11). \**p* < 0.05 compared to control by two-tailed unpaired t-test.

Another ion channel known to be regulated by NEDD4-2 is the amiloride-sensitive epithelial Na^+^ channel (ENaC), a constitutively active channel on the apical surface of high-resistance epithelial cells that mediates absorption of Na^+^ ions from the lumen in the distal nephron, distal colon, and the airways (*43*). ENaC-mediated Na^+^ absorption in the distal nephron regulates extracellular fluid (ECF) Na^+^ concentration, ECF volume and blood pressure, while ENaC in the lungs modulates airway surface liquid clearance (*43*). Structurally, ENaC is a heterotrimeric channel, comprised of membrane-spanning α, β, and γ subunits arranged in a counterclockwise orientation (*44*) (Fig. 4e). Physiologically, ENaC functional expression is regulated by NEDD4-2 which utilizes WW domains to bind PY motifs located on the carboxyl terminus of the β- and γ-subunits of the channel (*27*) (Fig. 4f). Mutations (missense, frameshift, or premature stops) that eliminate ENaC PY motifs lead to channel over-expression and result in Liddle syndrome, an autosomal dominant inherited form of severe and early-onset hypertension attendant with hypokalemia and metabolic alkalosis (*45–47*). HEK293 cells transiently transfected with ENaC α, YFP-tagged β, and γ subunits expressed constitutively-active amiloride-sensitive currents elicited by ramp and step protocols (Fig. 4g, h; Supplemental Fig. 11). Cells co-expressing WT α and γ with a Liddle syndrome mutant β-subunit, YFP-β[P617L] (*48*), displayed five-fold larger amiloride sensitive ENaC currents, consistent with the expected diminished restraint of channel surface density by endogenous NEDD4-2 (Fig. 4, f-h; Supplemental Fig. 11). Recruitment of endogenous NEDD4-2 to Liddle syndrome mutant ENaC (α, YFP-β[P617L], γ) using DiVa [nb.YFP-nb.C11] resulted in a marked decrease in amiloride-sensitive currents to levels that were not significantly different from control WT channels (Fig. 4g, h), demonstrating a reassertion of NEDD4-2 control of channel expression.

As a final test of the generality of the DiVa recruitment of endogenous NEDD4-2 approach, we returned to multi-subunit (membrane α_1B_, cytoplasmic Ca_V_β, and extracellular α_2_δ-1) Ca_V_2.2 channels. Reconstituted Ca_V_2.2 expressed with nb.F3 in HEK293 cells yielded robust whole-cell Ba^2+^ currents that were significantly inhibited by NEDD4-2 co-expression (Fig. 4i,j; Supplemental Fig. 12a,b). In accord, NEDD4-2 over-expression also decreased surface density of channels reconstituted with BBS-α_1B_ + β_2a_ + α_2_δ-1 subunits (Supplemental Fig. 12c,d). Co-expressing Ca_V_2.2 channel subunits with DiVa [nb.F3-nb.C11] resulted in a virtually identical decrease in whole-cell current amplitude and channel surface density as observed with NEDD4-2 over-expression (Fig. 4j; Supplemental Fig. 12). By contrast, DiVa [nb.F3-nb.C11] had no impact on BBS-Q1-YFP surface density (Supplemental Fig. 13), demonstrating substrate specificity of this approach. Finally, we tested the efficacy of recruitment of endogenous NEDD4-2 to inhibit native Ca_V_1/Ca_V_2 channels in their physiological context. We incorporated DiVa [nb.F3-nb.C11] into an adenovirus vector with tdTomato as a reporter, enabling expression in primary cultured murine dorsal root ganglion (DRG) neurons, a tissue that has been previously shown to express NEDD4-2 (Fig. 4k). Whole-cell patch clamp recordings of control cultured DRG neurons expressing just tdTomato yielded large high-voltage activated (HVA) Ca_V_ channel currents, reflecting the aggregate activity of all Ca_V_1/Ca_V_2 type channels present in these neurons (Fig. 4k,l). By contrast, DRG neurons expressing DiVa [nb.F3-nb.C11] expressed significantly reduced HVA Ca^2+^ currents (Fig. 4k,l), confirming the efficacy of recruiting endogenous NEDD4-2 to inhibit ion channels in their native environment.

### Comparative impact of NEDD4-2 over-expression and DiVa recruitment of endogenous NEDD4-2 on the global proteome

An important consideration related to whether, and to what extent, recruiting endogenous NEDD4-2 to a target substrate impacted global cellular proteostasis. To address this question, we utilized tandem mass tag (TMT) multiplexed mass spectrometry to conduct comparative total proteome analysis in three experimental groups comprised of HEK293 cells transiently transfected with Q1-YFP + nb.YFP-P2A-CFP (Group 1, control), Q1-YFP + NEDD4-2-P2A-CFP (Group 2) and Q1-YFP + DiVa [nb.YFP-nb.C11]-P2A-CFP (Group 3) (Fig. 5a). We used FACS to select CFP and YFP positive cells for processing to preclude trivial differences arising from variations in transient transfection efficiency (Fig. 5a). Reassuringly, NEDD4-2 was highly over-expressed in the proteome of cells from group 2 (transfected with NEDD4-2) relative to either group 1 (nb.YFP) or group 3 (DiVa[nb.YFP-nb.C11]), providing an internal positive control of the assay (Fig. 5b,d). Comparing the proteomes of group 1 to group 2 indicated NEDD4-2 over-expression resulted in 189 down-regulated proteins and 606 up-regulated proteins compared to control cells expressing nb.YFP (*p* = 0.05 and fold change (FC) = 1.5 cutoff) (Fig. 5b). By contrast, expression of DiVa [nb.YFP-nb.C11] resulted in a dramatically lower change in the cellular proteome (1 down-regulated and 17 up-regulated proteins) compared to control (Fig. 5c). Comparing groups 2 and 3 indicated NEDD4-2 over-expression resulted in 120 down-regulated proteins and 240 up-regulated proteins, respectively, compared to cells expressing DiVa [nb.YFP-nb.C11] (adj. *p* = 0.05 and FC = 1.5 cutoff) (Fig. 5d).

**Figure 5.**
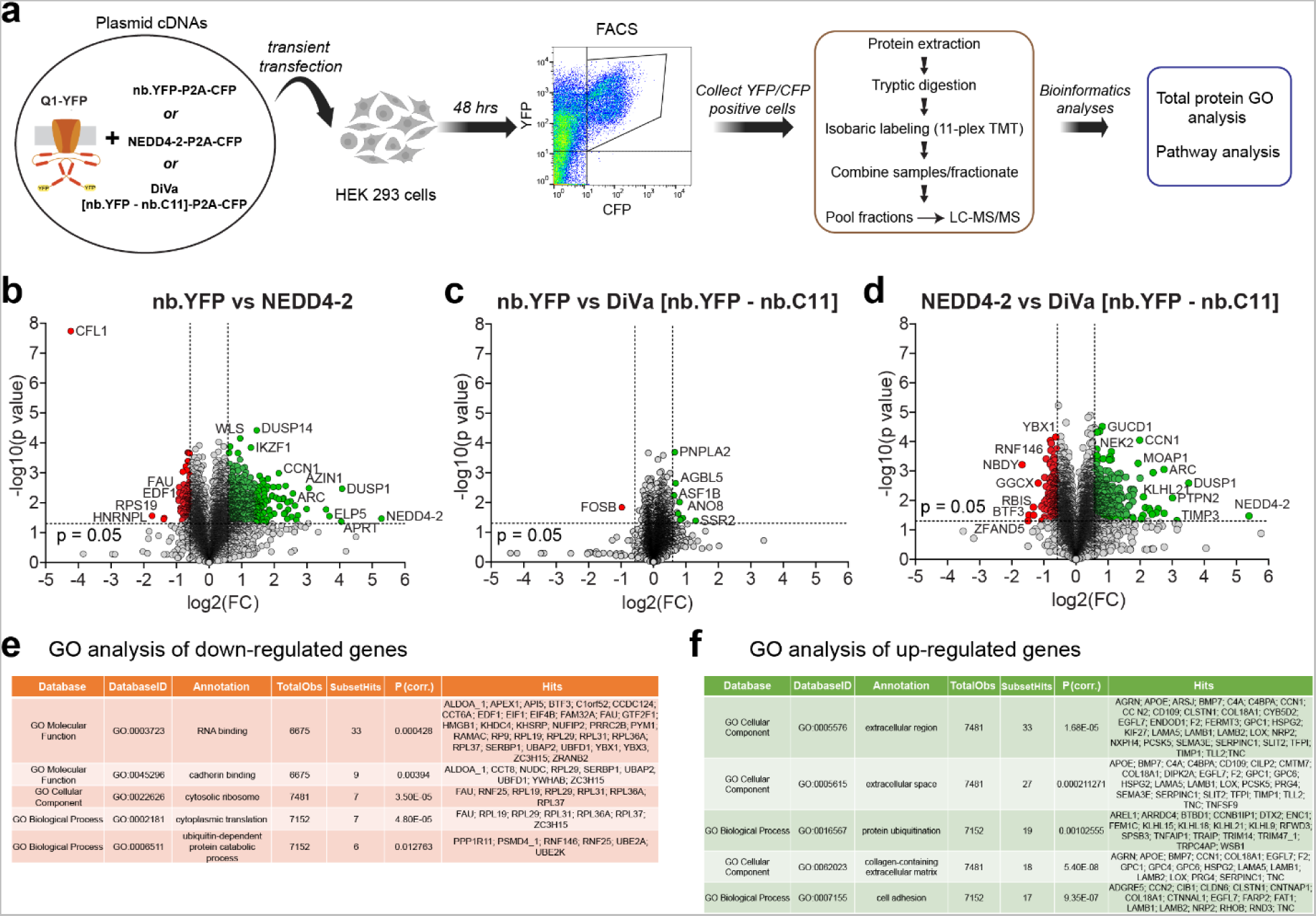
Comparative impact of NEDD4-2 over-expression and DiVa recruitment of endogenous NEDD4-2 on the proteome. (a) Schematic of experimental flow chart for tandem mass tag (TMT) mass spectrometry analyses of HEK293 cells proteome-wide changes. (b-d) Volcano plots comparing proteomes of HEK293 cells expressing (b) Q1-YFP + nb.YFP vs Q1-YFP + NEDD4-2, (c) Q1-YFP + nb.YFP vs Q1-YFP + DiVa [nb.YFP-nb.C11], and (d) Q1-YFP + DiVa [nb.YFP-nb.C11] vs Q1-YFP + NEDD4-2. Dotted lines demarcate fold-change (FC) cut-off = 1.5 and *p*-value = 0.05. Down-regulated genes fitting the cut-off criteria are colored in red and up-regulated genes in green. (e,f) Top terms from GO analyses of down-regulated and up-regulated genes, respectively, from (d).

Gene ontology (GO) analysis showed that genes down-regulated by NEDD4-2 over-expression were enriched in those involved in RNA binding or associated with ribosomes, with the top three biological processes potentially affected being protein translation, ubiquitin-dependent catabolism, and regulation of mRNA stability (Fig. 5e). Conversely, proteins that were up-regulated in NEDD4-2-over-expressing HEK293 cells were enriched for proteins in the extracellular space, the endoplasmic reticulum and Golgi lumen, the basement membrane, and the lysosomal lumen; the top three biological processes potentially affected include protein ubiquitination, cell adhesion, and cell migration (Figure 5f). Altogether, the data indicate that DiVa-mediated recruitment of endogenous NEDD4-2 to diverse ion channels achieves the same efficacy as NEDD4-2 over-expression on inhibition of ionic currents, but without the overt impact on proteostasis observed with NEDD4-2 over-expression.

## Discussion

Ion channels represent the second largest class of drug targets after GPCRs, and selective posttranslational ion channel blockers are highly sought after therapeutics for numerous diseases (*49*). Nevertheless, <10% of ion channels are targeted by currently approved medicines, and very few drugs in this category have been approved over the last three decades (*50*). A significant bottleneck is the identification of ligands that act as selective inhibitors of distinct ion channels. Typically, these are identified empirically by either high throughput screening and medicinal chemistry of small molecule libraries or isolated from complex mixtures of toxins that are present in venomous animals (*51*). These approaches are laborious, relatively slow, and often end up with ion channel blockers that are not selective for a particular channel isoform. Targeted protein degradation (TPD) offers a promising general approach to create new selective ion channel blockers. While there has been a widespread adoption of this technology to selectively eliminate proteins for either investigative or therapeutic purposes, the efficacy of the approach for composite ion channel protein complexes has not been previously demonstrated (*3*). This work definitively validates the general utility of creating novel ion channel inhibitors by targeted recruitment of an endogenous E3 ligase to a chosen ion channel.

The minimal requirements to successfully engage TPD boils down to identifying suitable binders for an endogenous E3 ligase and the target protein. An acknowledged deficiency in the TPD field is that <2% of the ~600 E3 ligases present in the human genome have been exploited for the method (*3*). We show here not only the utility of recruiting endogenous NEDD4-2 to diverse ion channels as being efficacious for their functional inhibition, but also the feasibility of using nanobodies for this purpose. This is the first demonstration that a HECT family E3 ligase is suitable for TPD which is notable because they have a unique mechanism of enzymatic action relative to the other E3 ligase families (*12, 52*). Moreover, their enzymatic activity is regulated by intricate intramolecular and posttranslational mechanisms that made it uncertain whether they would be suitable for TPD (*53*). NEDD4-2 was our strategic choice for this study because it has been demonstrated to physiologically regulate functional expression of many surface ion channels (*22, 54*). Beyond ion channels, NEDD4-2 also physiologically regulates the functional expression of diverse transporters and G-protein coupled receptors raising the prospect that it could be generally useful in TPD regulation of all classes of integral surface membrane proteins (*22*).

A recent proteome-wide screen to discover viable protein degradation and stabilization effectors did not identify any HECT E3 ligase family members as being suitable for TPD (*55*). The invisibility of HECT E3 ligases from this screen could be due to two main factors. First, multi-subunit ion channel complexes may be underrepresented in the choice of targets screened in this proteome-wide assay. Second, the sole focus on protein expression levels as a readout for efficacy may not be sufficient to identify effectors that are effective for down-regulation of ion channel function. Ion channels can be strongly inhibited by interventions that redistribute them from the plasma membrane to intracellular sites without their degradation (*29, 56*). The HECT domains of different NEDD4 family E3 ligases specify distinct polyubiquitin linkage types that can exert disparate functional outcomes on membrane protein degradation and trafficking (*57, 58*). The yeast homolog of NEDD4, Rsp5, preferentially catalyzes K63 polyubiquitin chains which are predominantly associated with non-degradative outcomes, including decreasing the surface density of target membrane proteins by promoting their net retention in intracellular sites (*22, 59, 60*). A pertinent question is how effective recruitment of endogenous NEDD4-2 would be in degrading cytosolic proteins which represent the largest class of proteins targeted by PROTACs molecules to date. While global proteomic analysis indicated NEDD4-2 over-expression results in decreased expression of many cytosolic and nuclear proteins, it is unclear which of these are due to a direct interaction with NEDD4-2, or are secondary to changes in proteins involved in mRNA stability, protein translation, or ubiquitin-dependent catabolism as identified by our GO analysis. Indeed, of 43 proteins identified to bind and/or be ubiquitinated by NEDD4-2 in an *in vitro* assay that were present in our proteome analysis (*61*), none were represented in the list of 120 down-regulated genes (NEDD4-2 vs. Diva [nb.YFP-nb.C11]), while one of these proteins (SENP2) was up-regulated. This could reflect either a lack of interaction between these proteins and NEDD4-2 in cells in contrast to observations *in vitro* or that the ubiquitination catalyzed by NEDD4-2 yields predominantly non-degradative outcomes. Yet another possibility is that another signaling event is required for over-expressed NEDD4-2 to catalyze degradation of particular protein substrates.

The results emphasize the versatility of nanobodies as a platform for developing selective binders for both E3 ligases and target proteins. In addition to their relatively small size (12-15 kDa) and ability to recognize antigens in the cytoplasm when expressed intracellularly, the ability of nb.C11 to distinguish between highly conserved NEDD4-1 and NEDD4-2 HECT domains highlights the exquisite selectivity of these reagents, which may not be as easily attainable using small molecules. Methods for identifying nanobodies specific for target proteins using immunization of alpacas or screening of synthetic nanobody display libraries are now readily available (*30, 62*). The advent of *in silico* de novo nanobody/antibody design and optimization algorithms (*63, 64*) may ultimately lower the barrier for developing these useful reagents, potentially allowing for generation of nanobodies against targets which are challenging to express and purify. The DiVa nanobody approach only requires identifying inert binders of the E3 ligase and the target protein. On the target protein side, this is a significantly lower bar than conventional approaches for identifying ion channel inhibitors. Overall, we envision that this work will enable a pipeline for high-throughput development of DiVa nanobody inhibitors for the entire surface ion channelome, with significant implications for both basic research and therapeutic development. Anticipated advantages of such DiVa nanobody inhibitors include the possibility of engineering tissue specificity by incorporating tissue-specific promoters and the potential to engineer chemogenetic or optogenetic variants that enable reversible control of channel inhibition by small molecules and light, respectively (*65*). Other induced proximity approaches specialized for eliminating surface membrane proteins have been developed and include LYTACS, AbTACS, PROTABs, and TransTACs (*18–20*). As part of their modes of action, all these approaches require generation of an extracellular antibody to the target membrane protein. The DiVa nanobody approach described here is fundamentally different to these prior methods in that both the target and E3 ligase effector are engaged intracellularly. The DiVa nanobodies are relatively small (<30 kDa) and fit comfortably within the packaging capacity of commonly used adeno-associated viral vectors. It may also be possible to pair a substrate-targeting nanobody with a ubiquitin variant that binds NEDD4-2 (or other HECT domain E3 ligase) without blocking catalysis to achieve targeted down regulation of protein expression (*28*). Altogether, the DiVa nanobody method offers the unique advantages of being well-suited for membrane proteins with large intracellular and small extracellular footprints for which it may be challenging to raise outward facing antibodies, and also being amenable for gene therapy development.

## Materials and Methods

### Molecular biology and plasmid construction

A full list of DNA constructs and related information for all DNA constructs used in this study is provided in the Excel document. In the following paragraphs, we describe cloning methods for constructs that were used for the first time in this study.

For NEDD4-2 HECT domain (or HECT4-2) purification, the HECT domain consisting of residues 615-994 of full-length NEDD4-2 (a gift from Joan Massague, Addgene plasmid # 27000) (*66*) was cloned via Gibson assembly into a NYCOMPS expression vector (a generous gift from Dr. Filippo Mancia) (*67*), with an N-terminal FLAG-10xHis-TEV cassette and a C-terminal FLAG tag.

Candidate HECT4-2 nanobodies were cloned into the PiggyBac CMV mammalian expression vector, and downstream of a cerulean marker using AgeI/NotI cloning sites. CFP-P2a-nb.C11 was generated by cloning nb.C11 into a previously described customized vector CFP-P2a-XX (*68*) using BamHI and NotI sites. HECT4-2 and HECT4-1 were both cloned into PCDNA3.1 and upstream of a Venus (Ven) marker using EcoRI/HindIII cloning sites. Full-length NEDD4-2 was cloned into PCDNA3.1-HECT4-2-Venus using BamHI/NheI to create NEDD4-2-Venus. To generate Venus-tagged HECT4-2 N- and C-lobes, the N- and C-lobes of NEDD4-2 HECT domain were cloned separately into a customized mammalian expression vector and downstream of mVenus using NotI and XbaI sites.

Human epithelial sodium channel (ENaC) α, β, and γ subunits were gifts from Christie Thomas (Addgene plasmid # 83430, 83429, and 83429, respectively) (*69*). To make Venus-β-ENaC, mVenus was cloned via BamHI/KpnI cloning sites into the human β-ENaC construct. Ven-β-ENaC P617L was created using PCR mutagenesis.

### Cell culture and transfection

Low passage HEK293 cells were cultured in DMEM supplemented with 8% fetal bovine serum (FBS) and 100 mg/mL of penicillin–streptomycin and maintained in humidified incubators at 37°C and 5% CO_2_. HEK293 cell transfection was accomplished using the calcium phosphate precipitation method. Briefly, for transfections performed in 12-well plates, 1 µg of each plasmid to be transfected was mixed with 7.75 μL of 2 M CaCl_2_ and sterile deionized water (to a final volume of 62μL). The mixture was then added dropwise, with constant tapping, to 62 μL of 2x HEPES buffered saline containing (in mM): HEPES 50, NaCl 280, Na_2_HPO4 1.5, pH 7.09. The resulting DNA–calcium phosphate mixture was incubated for 20 min at room temperature (RT) and then added dropwise to HEK293 cells (60 – 80% confluent). Cells were washed with Ca^2+^-free phosphate buffered saline after 4-6 h and maintained in supplemented DMEM. For other applications, the volume and amounts of transfection reagents were appropriately scaled based on the area of the transfection culture dish.

Chinese hamster ovary (CHO) cells were obtained from ATCC (Manassas, VA), and cultured at 37°C in Kaighn’s Modified Ham’s F-12K (ATCC) supplemented with 8% FBS and 100 mg/mL of penicillin–streptomycin. CHO cells were transfected using the Lipofectamine™ 3000 Transfection Reagent, following the manufacturer’s instructions.

### DRG neuron isolation and culture

The isolation of murine dorsal root ganglion (DRG) neurons was performed following the Columbia University Institutional Animal Care and Use Committee guidelines. Adult WT C57BL mice were used in this study. Following decapitation of mice, DRGs were harvested and transferred to Ca^2+^-free, Mg^2+^-free Hanks’ balanced salt solution (HBSS; Thermo Fisher Scientific #14170161). Ganglia were treated with collagenase P (1.5 mg/mL; Sigma-Aldrich, #11249002001) in HBSS for 20 min at 37 °C (with gentle shaking), followed by a 2-minute incubation at 37 °C (with gentle rotation) in 1 mL warm TrypLE (Thermo Fisher Scientific, #12604013). The majority of TrypLE was removed by pipetting after incubation. Leftover TrypLE was neutralized with culture medium (modified Eagle’s medium [MEM]; Thermo Fisher Scientific, #11095-080) supplemented with 10% horse serum (heat-inactivated; Thermo Fisher Scientific #26050070), 100 U/mL penicillin, 100 μg/mL streptomycin (Thermo Fisher Scientific, #15140122), 1% MEM vitamin solution (Thermo Fisher Scientific, #11120052), and 2% B-27 supplement (Thermo Fisher Scientific, #17504044). The DRGs were left in the tubes vertically for 5 min after which the serum-containing medium was decanted. DRGs were then resuspended with serum-free MEM containing the supplements listed above and triturated using a fire-polished Pasteur pipette. DRG neurons were plated on laminin-treated (0.05 mg/mL) glass coverslips previously rinsed with 70% ethanol and sterilized by UV light exposure for at least 15 minutes. DRG cultures were then incubated at 37 °C in 5% CO_2_ for ~3-6 hrs, after which adenoviral vectors (mCherry control; tdTomato-P2a-DiVa [nb.β-nb.C11]; Vector Biolabs (Malvern, PA) were added separately to respective DRG culture wells. The DRG cultures were then maintained at 37 °C in a 5% CO_2_ humidified incubator and used for electrophysiological experiments at ~48 hrs post-adenoviral infection.

### Flow cytometry-based cell surface labelling assay

Cell surface ion channel pools were assayed by flow cytometry in live, transfected HEK293 cells as previously described (*29*). 48 hours post-transfection, cells were gently washed with ice-cold phosphate-buffered saline (PBS) containing Ca^2+^ and Mg^2+^ (in mM: 0.9 CaCl_2_, 0.49 MgCl_2_, pH 7.4), and then incubated for 30 min in blocking medium (DMEM with 3% BSA) at 4°C. Afterward, the HEK293 cells were incubated with 1 μM Alexa Fluor-647 conjugated α-bungarotoxin (BTX-647; Life Technologies) at 4°C for 1 hr. Subsequently, the HEK293 cells were washed three times with PBS containing Ca^2+^ and Mg^2+^. HEK293 cells were then gently harvested in Ca^2+^-free PBS, and assayed by flow cytometry using either a BD Fortessa or BD LRSII Cell Analyzer (BD Biosciences, San Jose, CA, USA). CFP, YFP, and mCherry-tagged proteins were excited at 405 nm, 488 nm, and 561 nm, respectively, while Alexa Fluor-647 was excited at 633 nm (on the LRSII) or 640 nm (on the Fortessa).

### Electrophysiology

For voltage-gated calcium channel and epithelial sodium channel (ENaC) electrophysiology experiments, whole-cell recordings from HEK293 cells and DRG neurons were conducted at room temperature 48 hrs after transfection using EPC-8 and EPC-10 patch-clamp amplifiers (HEKA Electronics) controlled by pulse software (HEKA). Micropipettes were prepared from 1.5 mm thin-walled glass (World Precision Instruments) using a P97 microelectrode puller (Sutter Instruments). For calcium channel recordings in both HEK293 cells and DRG neurons, the internal solution contained (in mM): 135 cesium methansulfonate (CsMeSO_3_), 5 CsCl, 5 EGTA, 1 MgCl_2_, 10 HEPES, and 2 mg/mL MgATP (pH 7.3). Series resistance was typically between 1-2 MΩ. There was no electronic resistance compensation. The external solution contained (in mM): 140 tetraethylammonium-MeSO_3_, 5 BaCl_2_, and 10 HEPES (pH 7.4). Whole-cell I-V curves were generated from a family of 20 ms-step depolarizations (−40 mV to +120 mV from an initial 10 ms hold at −90 mV). There was a final repolarization step to −90 mV for 20 ms. Currents were sampled at 20kHz and filtered at 5kHz. Traces were acquired at a repetition interval of 10s. Leak and capacitive transients were subtracted using a P/4 protocol.

Whole-cell measurements of amiloride-sensitive currents were performed in HEK293 cells transfected with the alpha, YFP-tagged beta, and gamma subunits of the epithelial sodium channel (ENaC). The pipette solution was composed of (in mM): 150 KCl, 2 MgCl_2_, 5 EGTA, and 10 HEPES. The external solution contained (mM): 150 NaCl, 2 MgCl_2_, 2 CaCl_2_, 10 HEPES, and 0.01 amiloride. The initial test external solution did not contain amiloride. The current due to ENaC was defined as the amiloride-sensitive current, which is the difference between currents obtained in the absence and presence of 0.01 mM amiloride. Series resistance was typically between 2-3 MΩ. There was no electronic resistance compensation. To define the amiloride-sensitive ENaC currents, a voltage-ramp protocol was first applied to transfected HEK293 cells with the cells initially in an external solution without amiloride. For this voltage-ramp protocol, transfected HEK293 cells were held at +40 mV for 20 ms; followed by a step to +60 mV for 5 ms; then a 500 ms voltage-ramp to −100 mV; and a return to +40 mV holding potential for 20 ms. Currents were sampled at an interval of 1 ms and filtered at 500 Hz with a 2.5 s interval between successive sweeps. After currents became stable, a voltage-step protocol was applied with the cells in the external solution without amiloride. For the voltage-step protocol, the cells were held at +40 mV for 20 ms followed by a family of 500 ms steps from −120 mV to +80 mV in 20-mV increments, and a return to +40 mV holding potential for 20 ms. Currents were sampled at 1 kHz and filtered at 333 Hz with a 2.5 s sweep interval. After the initial recordings in the external solution without amiloride, the voltage-ramp and voltage-step protocols were repeated in the named order for the same cell a second time while perfusing an external solution containing 0.01 mM amiloride. A second voltage step protocol was then conducted in the presence of amiloride.

For electrophysiology recordings of KCNQ1 currents, transfected HEK293 and CHO cells were plated on 3.5 cm culture dishes on the stage of an inverted microscope (OLYMPUS BH2-HLSH, Precision Micro Inc, Massapequa, NY, USA). The external solution contained the following: 132 mM NaCl, 4.8 mM KCl, 2 mM CaCl_2_, 1.2 mM MgCl_2_, 10 mM HEPES, and 5 mM glucose (pH was adjusted to 7.4 with NaOH). The internal solution contained the following: 110 mM KCl, 5 mM ATP-K_2_, 11 mM EGTA, 10 mM HEPES, 1 mM CaCl_2_, and 1 mM MgCl_2_ (pH was adjusted to 7.3 with KOH). Pipette series resistance was typically 1.5–3 MΩ when filled with the internal solution. Whole-cell currents were recorded at room temperature using an Axopatch 200B amplifier (Axon Instruments, Foster City, CA, USA). Current-voltage relationship curves were generated using 2-second depolarizing steps ranging from −60 mV to +100 mV (in 20 mV increments). The currents were triggered every 10 seconds from an initial 500 ms hold at −70 mV, and a final repolarizing step to −40 mV for 5 seconds. Currents were sampled at 10 kHz and filtered at 5 kHz.

### Flow cytometry FRET (Flow-FRET) experiments

The experimental and analysis steps used for the flow cytometry-based fluorescence resonance energy transfer (Flow-FRET) assay have been previously described in detail (*31, 68, 70*). Briefly, HEK293 cells were transfected using the calcium phosphate transfection method (described above) or the polyethylenimine (PEI) transfection method. 1 μg each of cerulean (Cer)- and Venus (Ven)-tagged cDNA pair was mixed in 100 μL of serum-free DMEM media. Afterward, 5 μL of PEI was added to each experimental condition. The DNA-PEI mixture was allowed to sit at room temperature for 15-25 minutes following which the mixture was added to the cells. FRET experiments were performed ~48 hrs post-transfection. 100 μM cycloheximide was added to cells ≥2 hrs before experimentation to halt the synthesis of new fluorophores and to allow existing fluorophores to fully mature. Cells were gently washed with ice-cold PBS (containing Ca^2+^ and Mg^2+^), harvested in Ca^2+^-free PBS, and assayed by flow cytometry using a BD LSR II Cell Analyzer (BD Biosciences). Cerulean, Venus, and FRET signals were analyzed using the following laser and filter set configurations: BV421 (excitation, 405 nm; emission, 450/50 nm), FITC (excitation, 488 nm; emission, 525/50 nm) and BV520 (excitation, 405 nm; emission, 525/50 nm). Several controls were prepared for each experiment, including untransfected blanks for background subtraction, single-colour Venus and Cerulean for spectral unmixing, Cerulean and Venus co-expressed together for concentration-dependent spurious FRET estimation, as well as a series of Cerulean–Venus dimers for FRET calibration. Custom MATLAB software was used to analyze FRET donor and acceptor efficiency and generate FRET binding curves as a function of the concentrations of free acceptor and donor.

### Protein purification for nanobody generation

For NEDD4-2 HECT domain purification, the NEDD4-2 HECT domain plasmid was transformed into Rosetta DE3 *E. coli* (Millipore Sigma), following manufacturers’ instructions. Cells were grown at 37°C in 1L 2xTY media supplemented with 50 μg/mL carbenicillin and 35 μg/mL chloramphenicol and shaken at 225 rpm. Protein expression was induced with 0.2 mM IPTG when the cells reached an OD of 0.6-0.8. The cells were then grown overnight at 22°C. Cells were lysed using an Avestin® Emulsiflex-C3 homogenizer in a buffer containing 50 mM Tris, 150 mM KCl, 10% sucrose, 1 mM PMSF (phenylmethylsulfonyl fluoride), and EDTA-free cOmplete protease inhibitor cocktail (Roche), pH 7.4. The lysate was spun down at 35,000g for 1 hr. NEDD4-2 HECT domain was subsequently isolated from the supernatant with anti-FLAG antibody (M2) affinity chromatography and eluted with 100 μg/mL FLAG peptide (Sigma Millipore) in 50 mM TrisHCl, 150 mM KCl, pH 7.4. The protein was then applied to an ion exchange column (MonoQ, GE) and eluted with a linear KCl gradient of 50 mM to 1M. Peak fractions were collected and subjected to size exclusion chromatography (Superdex 200, GE) in a buffer containing 20 mM Tris, 150 mM KCl, pH 7.4. Proteins were brought to 20% glycerol, flash frozen, and stored at −80°C.

### HECT4-2 nanobody (nb.C11) isolation from yeast nanobody display

We used a synthetic yeast nanobody display library (kind gift from Dr. Andrew Kruse, Harvard University) to isolate nanobody binders to purified HECT4-2 using a previously described protocol (*30*). The naive yeast library was initially grown in 250 mL sterilized ‘Trp/Gal’ media, composed of the following: 0.48 g Trp drop-out media supplement (Sigma Aldrich), 1.675 yeast nitrogen base, 2.6 g sodium citrate, 1.85 g citric acid monohydrate, 2.5 mL Pen/Strep (10,000 units/mL stock), and 5 g galactose, pH 4.5. Cells were incubated for 48 hours at 25°C, 220 rpm to induce nanobody expression. Induced cells were washed and incubated in the following buffer for selection: PBS, 0.1% BSA, and 5 mM maltose. For the first selection, 5×10^9^ cells were first subject to a preclear step, to remove nanobodies that bind selection reagents. Cells were incubated with 250 μL anti-FITC microbeads (Miltenyi) and 75 μL FITC-conjugated anti-FLAG M2 antibody (Sigma Aldrich) at 4°C for 30 minutes, and then passed through an LD column. These pre-cleared cells were then subjected to a positive selection by incubating the cells with 1 μM HECT4-2 and the same amount of microbeads and anti-FLAG FITC, at 4°C for 1 hr and then passed through an LS column. The enriched population of yeast was quantified (to approximate the library size) and then grown for 48 hours at 30°C, 220 rpm. Subsequent selections used 10x the number of cells obtained after the previous selection, and 1 μM HECT4-2. After two rounds of MACS selection, the yeast were stained with 1 μM HECT4-2 and sorted into single wells of a 96-well plate using fluorescence-activated cell sorting (FACS). Individual clones were grown and used for binding validation and subsequent plasmid isolation.

#### Expression and purification of WW4-HECT domain

The WW4-HECT domain (residues 560-994 of Addgene #2700), corresponding to residues 541-975 in the canonical full-length human NEDD4-2 sequence (Uniprot id: Q96PU5-1) was cloned into NYCOMPS expression plasmid (*67, 71*) via the Gibson assembly method (*72*) with N-terminal Flag-His_10_ tags followed by a TEV cleavage site (ENLYFQSY) and a C-terminal Flag tag (Fig 3a, right). The plasmid containing WW4-HECT (NEDD4-2) was transformed into *E. coli* BL21 (DE3) pLysS. Cells were grown at 37°C, shaken at 280 rpm in 1 L Terrific Broth supplemented with 50 μg/ml Carbenicillin, 35 μg/ml chloramphenicol and 0.4% glycerol until OD600 0.5, whereupon expression was induced by the addition of 1 mM IPTG, and cells were grown at 18 °C for a further 18-20 hours. The cells were harvested by centrifugation at 10,000 rpm for 10 mins at 4°C using Beckman Coulter JLA-10.500 Fixed Angle rotor, resuspended in 1x Phosphate Buffered Saline to remove excess media, centrifuged again and stored at −80°C prior until use. Thawed cell pellets were resuspended in His Equilibration buffer (HE) containing 20 mM Tris HCl pH 7.5, 300 mM NaCl, 10% glycerol supplemented with 0.5 mM PMSF, 1 mM TCEP, 1 mM EDTA, 10 μg/mL DNase I and EDTA-free cOmplete^TM^ protease inhibitor cocktail (Roche) and lysed using Avestin® Emulsiflex-C3 homogenizer. Lysates were clarified by ultracentrifugation at 34,000 rpm for 45 minutes at 4°C in a Type 45.1 Ti rotor (Beckman-Coulter). The supernatants were then passed through a 0.22 μm syringe sterile filter (Thermo-Fisher) before being applied to 10 mL of Ni-NTA XPure Agarose Resin (UBPBio) pre-equilibrated with HE buffer without EDTA and with 20 mM Imidazole. After the resin was extensively washed with 10 column volumes of wash buffer (HE + 70 mM imidazole, pH 7.5), the WW4-HECT protein was eluted with 2-4 column volumes of elution buffer (HE + 350 mM imidazole, pH 7.5). The eluate buffer was exchanged into a buffer containing 20 mM HEPES pH 7.5, 200 mM NaCl, and 10% glycerol using a desalting column to remove imidazole. WW4-HECT protein was treated with recombinant TEV protease (molar ratio 25:1) for 3 hours at room temperature to cleave the N-terminal Flag and His tags. Uncleaved protein and cleaved tag peptide was captured by passing over a 1 mL Ni-NTA XPure Agarose Resin (UBPBio). The cleaved WW4-HECT protein was concentrated (Amicon Ultra-15 centrifugal filter, 10 kDa cutoff; EMD Millipore), filtered, and further purified by size-exclusion chromatography on a Superdex 200 Increase 10/300 GL column (Cytiva) in 20 mM HEPES pH 7.5, 150 mM NaCl, 1mM TCEP and 1 mM EDTA. The peak fractions containing WW4-HECT domain were evaluated for purity by SDS-PAGE, pooled and concentrated to yield 25 mg purified protein per liter of cell culture as evaluated by absorbance at 280 nm, and flash frozen for future use.

#### nb.C11 Expression and purification

Nb.C11 was cloned into a periplasmic expression vector (NYCOMPS) containing an N-terminal pelB leader sequence and a C-terminal hexahistidine tag and expressed in 2 L cultures of *E. coli* BL21 (DE3) pLysS strain. Expressed nb.C11 was nickel-affinity purified by a previously described protocol (*30*). The nanobody eluted from 5 mL Ni Sepharose 6 Fast Flow resin (Cytiva) in 20 mM HEPES pH7.5, 100 mM NaCl supplemented with 400 mM Imidazole was concentrated to 400 μl and further purified by size-exclusion chromatography on a Superdex 200 Increase 10/300 GL column (Cytiva) in 20 mM HEPES pH 7.5, 150 mM NaCl, 5% glycerol. The peak fractions of the nb.C11 were pooled, concentrated and flash frozen for future use.

#### WW4-HECT:nb.C11 complex formation and cryo-EM specimen preparation

Size-exclusion chromatography fractions of purified WW4-HECT protein were mixed with nanobody at a molar ratio of 1:1.5 and incubated for 30 mins on ice. The mixture was subjected to size-exclusion chromatography on a Superdex 200 Increase 10/300 GL column in a buffer containing 20 mM HEPES pH 7.5, 150 mM NaCl, 1 mM TCEP, and 1 mM EDTA. The fractions corresponding to WW4-HECT:nb.C11 complex were validated for purity and the presence of both components by SDS-PAGE. The peak fractions of the complex were then concentrated to ~11 mg/mL ((157 μM). The complex was further subjected to mass photometry (Refeyn OneMP) analysis which showed a sharp, 69-kDa molecular weight peak corresponding to the expected molecular weight of the WW4-HECT:nb.C11 complex with a 17 kDa shift from measurement of the WW4-HECT alone. Right before plunge cooling, 1 μL of 0.3% CHAPS was added to a 6 μL sample of WW4-HECT:nb.C11 complex and homogenously mixed by pipetting so that the final protein and CHAPS concentrations were at 9 mg/mL and 0.043% respectively. 3 μL of the WW4-HECT:nb.C11 complex was applied to the foil side of glow-discharged UltraAuFoil 0.6/1µm holey gold grids (SPT Labtech), and blotted using 595 filter paper (Ted Pella, Inc) for 9s with a blot force of 4 and a wait time of 30 sec using a Vitrobot Mark IV (Thermo Fisher Scientific) operated at 4°C and 100% humidity. Each grid was vitrified by plunging into liquid ethane precooled in liquid N_2_.

#### EM Data Acquistion

Cryo-EM data for the WW4-HECT:nb.C11 complex were recorded on a Titan Krios electron microscope (Thermo Fischer Scientific) operated at 300 kV, equipped with Gatan K3 Bio-Quantum with a silt width of 20 eV and a Gatan K3 direct electron detector. The energy filter slit was aligned automatically every hour using Leginon. All the cryo-EM movies were recorded in counting mode and with a nominal magnification of 165,000x which corresponds to a pixel size of 0.5135 Å, calibrated by crystal structure correlation using an apoferritin dataset collected under the same conditions. The data collection was performed using a dose of ~109.2 e^-^/Å^2^ (*73–75*) across 100 frames at a dose rate of approximately 16.1 e^-^/pix/s, corresponding to dose per frame of 1.092 e^-^/Å^2^ and using a set defocus range of −1 to −2.5 μm. A 100 µm objective aperture was used. In total, 7,790 micrographs were recorded over a two-day collection.

#### EM Data processing

CryoSPARC v4.4 was used for all processing steps unless otherwise indicated. 7,790 movies were imported into CryoSPARC, and dose-weighted and motion corrected with Patch Motion. Patch-based CTF estimation was performed using Patch CTF on the aligned, non-dose-weighted averages followed by manual curation of micrographs, leaving 5,742 micrographs for further processing. Blob picker was used to pick particles with 100 Å and 200 Å minimum and maximum particle diameter respectively. 1,631,714 particles were extracted and downsampled from 512 to 200-pixel box size. The particles were subjected to two rounds of 2D classification with 80 online-EM iterations and 5 full iterations. 2D classes with well-defined high-resolution features were selected, leaving 116,543 particles. *Ab initio* reconstruction was performed using 4 classes, with default settings. This resulted in one unique class with the nb.C11 bound to the HECT domain in the T-state conformation, and three less well-defined classes apparently corresponding to the HECT domain alone. Particles from the four *ab initio* were used as input for heterogeneous refinement, with default settings. The nanobody-bound class from heterogeneous refinement was subjected to non-uniform refinement, yielding a final resolution of 3.95 Å (23,078 particles). Particles were extracted with recentering while downsampling from 512 to 200-pixel box size as before, followed by non-uniform refinement, including on-the-fly refinement of til/trefoil and global refinement of beam tilt and trefoil aberrations yielding an initial consensus refinement resolution of 3.74 Å (22, 904 particles). A Topaz model was trained (*76*) using these 22, 904 particles, with a downsampling factor of 8, and an estimated number of particles per micrograph of 50. The resulting model was used to pick particles using Topaz Extract. Heterogenous refinement was performed on the 742,103 particles picked with Topaz using the four ab initio models and four random density decoys, resulting in identification of one good nanobody-bound class of 152,408 particles, which was refined to 3.16Å using non-uniform refinement. Refinement after splitting into image shift groups further improved resolution to 3.11 Å. We performed 3D classification without alignments on the 152,408 particles using a focused mask around the nanobody, allowing identification of a class with improved density quality for the nanobody. Non-uniform refinement of this class resulted in the final 3.06 Å reconstruction from 90,979 particles. FSC validation was calculated with a mask around the nanobody and another mask around the WW4-HECT domain with a 12 pix soft edge (Supplemental Fig. 7) which allowed for calculating the overall resolution in the nanobody and the HECT domain to be 3.10 Å and 3.16 Å respectively. We then ran orientation diagnostics analysis in CryoSPARC 4.4 on the final reconstruction to analyze the orientation distribution and directional anisotropy of the data, which showed minimal anisotropy (Supplemental Fig. 4d) (*77, 78*). Detailed data collection, refinement and validation statistics are provided in Table 1.

#### Atomic Model Building and Refinement

We used the NEDD4-1 HECT domain crystal structure (PDB: 4BE8) as an initial model and placed it in the corresponding non-uniform refined and sharpened map and fit as rigid body in UCSF Chimera (*79*). For the nanobody, we used a ColabFold predicted structure of the nanobody C11 (*80*) as a model and fit as rigid body in UCSF chimera. For the WW4 domain, a previously published NMR structure of NEDD4-2 WW3 domain (PDB: 2MPT) (*33*) was used as an initial model, rigid-body fit in UCSF Chimera, and after combining the three parts of the atomic model, the complete assembly was completed and adjusted by real-space fitting in Coot (*81, 82*). Because the EM map did not exhibit any density for the C-terminus tail of HECT domain, and the residues 581-587 of the WW4 domain, these parts were truncated for the final maps of the WW4-HECT:nb.C11 complex. Finally, iterative rounds of real space-refinement of the whole complex were performed in PHENIX v1.21-5207. The atomic model was validated using Molprobity (*83*). A complete summary of data collection and refinement statistics are provided in supplemental Table 1.

### Immunoprecipitation and immunoblotting

Transfected HEK293 cells cultured in 60 mm dishes were harvested in PBS and then centrifuged at 2,000 g (4°C) for 5 min. The cell pellets were resuspended in RIPA lysis buffer containing (mM): 150 NaCl, 50 Tris HCl, 1 EDTA, 0.1% (w/v) SDS, 1% Triton X-100, 0.5% sodium deoxycholate, and supplemented with protease inhibitor mixture (10 μL/mL, Sigma Aldrich), 1 PMSF, 2 N-ethylmaleimide, 0.05 PR-619 deubiquitinase inhibitor (LifeSensors). Cells were lysed on ice for 1 hr with intermittent vortexing and then centrifuged at 10,000 x g for 10 min at 4°C. The cell lysate was collected following which protein concentration was determined with the bicinchoninic acid (BCA) protein estimation kit (Pierce Technologies).

For KCNQ1 pulldowns, cell lysates were precleared with 10 μL of protein A/G Sepharose beads (Rockland) for 1 hr at 4°C and then incubated with 1 μg anti-KCNQ1 antibody (Alomone Labs, Israel) for 1 hr at 4°C. Equivalent amounts of protein were then added to spin columns with 25 μL equilibrated protein A/G Sepharose beads and rotated overnight at 4°C. Immunoprecipitates were washed five times with RIPA buffer and then eluted with 40 μL elution buffer (50 mM Tris, 10% (vol/vol) glycerol, 2% SDS, 100 mM DTT, and 0.2 mg/mL bromophenol blue) at 55°C for 15 min. Proteins were resolved on a 4-12% Bis-Tris gradient precast gel (Life Technologies) in MOPS-SDS running buffer (Life Technologies) at 200V constant for ~1 h. Protein bands were transferred by tank transfer onto a polyvinylidene difluoride (PVDF, EMD Millipore) membrane in transfer buffer (25 mM Tris pH 8.3, 192 mM glycine, 15% (vol/vol) methanol, and 0.1% SDS). The membranes were blocked with a solution of 5% non-fat dry milk (BioRad) in tris-buffered saline-Tween20 (TBS-T) (25 mM Tris pH 7.4, 150 mM NaCl, and 0.1% Tween-20) for 1hr at RT and then incubated overnight at 4°C with primary anti-KCNQ1 (Alomone Labs) antibody in blocking solution. The blots were washed with TBS-T three times for 10 min per wash and then incubated with a goat anti-rabbit secondary horseradish peroxidase-conjugated antibody (Thermo Fisher Scientific) for 1 hr at RT. After washing in TBS-T, the blots were developed with a chemiluminescent detection kit (Pierce Technologies) and then visualized on a gel imager. Membranes were then stripped with harsh stripping buffer (2% SDS, 62mM Tris pH 6.8, 0.8% ß-mercaptoethanol) at 50°C for 30 min, rinsed under running water for 2 min, and washed with TBST (3 times, 10 min per wash). Membranes were pre-treated with 0.5% glutaraldehyde and re-blotted with anti-ubiquitin (VU1, LifeSensors) according to the manufacturers’ instructions.

For immuno-detection of total protein fractions, equal amounts of proteins from each treatment group were loaded. Sodium dodecyl sulfate-polyacrylamide gel electrophoresis (SDS-PAGE), gel transfer, and immunoblotting were performed as described above, but with appropriate antibodies for the detection of KCNQ1 and actin.

#### Confocal Microscopy

For confocal microscopy studies, freshly isolated murine DRG neurons were cultured on 12 mm diameter glass coverslips. The glass coverslips were placed within 35 mm tissue culture plates containing DRG culture media (described under the DRG culture section). Adenoviral infection of the cultured DRG neurons was done as described in the tissue culture section above. 5 days post-infection, the cultured DRG neurons were washed once in PBS and then fixed with 4% paraformaldehyde at room temperature (RT) for 15 minutes. The fixed DRG neurons were subsequently washed three times with 0.1 M glycine in PBS following which the glass coverslips containing the fixed DRG neurons were placed on glass microscope slides and mounted in 4′,6-diamidino-2-phenylindole (DAPI) stain. Images were captured with a Nikon Eclipse Ti2 inverted microscope equipped with 405 nm, 488 nm, 561 nm, 638 nm lasers and run by NIS-Elements AR 5.02.00 64-bit software. Images were detected using the Nikon PLAN APO λD 20X/0.80 objective and analyzed using Fiji software.

### Florescence-activated cell sorting (FACS)

For fluorescence-activated cell sorting (FACS), low passage HEK293 cells were cultured on 10 cm tissue culture plates and grown in DMEM supplemented with 8% fetal bovine serum (FBS) and 100 mg/mL of penicillin–streptomycin. The cells were maintained in humidified incubators at 37°C and 5% CO_2_. HEK293 cells were transiently co-transfected with YFP-tagged KCNQ1 (10 µg) and one of the following: CFP-P2a-nb.YFP (10 µg); or CFP-P2a-NEDD4-2 (10 µg); or CFP-P2a-DiVa[nb.YFP-nb.C11] (10 µg). FACS was done ~48 hrs post-transfection. In preparation for FACS, adherent HEK293 cells were dissociated from the respective cell culture plates using TrypLE; re-suspended in 2% heat-inactivated FBS in phosphate-buffered saline (PBS); passed through separate 40 μm filters; and kept on ice before sorting. FACS was carried out on either a BD FACSAria II or BD Influx cell sorter each equipped with 405 nm, 488 nm, 561 nm, and 638 nm excitation lasers. For cell sorting, untransfected, YFP-only expressing, and CFP-only expressing HEK293 cells were used to define the region of interest. HEK293 cells expressing both YFP (positive for KCNQ1-YFP) and CFP (positive for anti-YFP-nanobody, NEDD4-2 or DiVa) were collected after sorting. Approximately 1 million cells were sorted for each group with two to three biological replicates. Immediately following sorting, the sorted cells were re-suspended in 10% FBS in PBS and kept on ice. Afterward, the cells were washed twice in PBS by centrifugation to remove residual FBS. The cell pellets were stored at −80°C before and during transport for mass spectrometry studies.

### Sample Preparation for Mass Spectrometry

Samples for protein analysis were prepared essentially as previously described (*84, 85*). Proteomes were extracted using a buffer containing 200 mM EPPS pH 8.5, 8 M urea, 0.1% SDS and protease inhibitors. Following lysis, each sample was reduced with 5 mM TCEP. Cysteine residues were alkylated using 10 mM iodoacetimide for 20 minutes at RT in the dark. Excess iodoacetimide was quenched with 10 mM DTT. A buffer exchange was carried out using a modified SP3 protocol (*86*). Briefly, ~250 µg of Cytiva SpeedBead Magnetic Carboxylate Modified Particles (65152105050250 and 4515210505250), mixed at a 1:1 ratio, were added to each sample. 100% ethanol was added to each sample to achieve a final ethanol concentration of at least 50%. Samples were incubated with gentle shaking for 15 mins. Samples were washed three times with 80% ethanol. Protein was eluted from SP3 beads using 200 mM EPPS pH 8.5 containing trypsin (ThermoFisher Scientific, 90305R20) and Lys-C (Wako, 129-02541). Samples were digested overnight at 37°C with vigorous shaking. Acetonitrile was added to each sample to achieve a final concentration of ~33%. Each sample was labeled, in the presence of SP3 beads, with ~60 µg of TMT 11plex reagents (ThermoFisher Scientific). Following confirmation of satisfactory labeling (>97%), excess TMT was quenched by the addition of hydroxylamine to a final concentration of 0.3%. The full volume from each sample was pooled and acetonitrile was removed by vacuum centrifugation for 1 hour. The pooled sample was acidified using formic acid and peptides were de-salted using a Sep-Pak 50 mg tC18 cartridge (Waters). Peptides were eluted in 70% acetonitrile, 1% formic acid and dried by vacuum centrifugation.

### Basic pH reversed-phase separation (BPRP)

TMT labeled peptides were solubilized in 5% ACN/10 mM ammonium bicarbonate, pH 8.0 and ~300 µg of TMT labeled peptides were separated by an Agilent 300 Extend C18 column (3.5μm particles, 4.6 mm ID and 250 mm in length). An Agilent 1260 binary pump coupled with a photodiode array (PDA) detector (Thermo Scientific) was used to separate the peptides. A 45-minute linear gradient from 10% to 40% acetonitrile in 10 mM ammonium bicarbonate pH 8.0 (flow rate of 0.6 mL/min) separated the peptide mixtures into a total of 96 fractions (36 seconds). A total of 96 Fractions were consolidated into 24 samples in a checkerboard fashion and vacuum-dried to completion. Each sample was desalted via Stage Tips and re-dissolved in 5% FA/ 5% ACN for LC-MS3 analysis.

### Liquid chromatography separation and tandem mass spectrometry (LC-MS3)

Proteome data were collected on an Orbitrap Fusion Lumos mass spectrometer (ThermoFisher Scientific) coupled to a Proxeon EASY-nLC 1000 LC pump (ThermoFisher Scientific). Fractionated peptides were separated using a 120 min gradient at 550 nL/min on a 35 cm column (i.d. 100 μm, Accucore, 2.6 μm, 150 Å) packed in-house. MS1 data were collected in the Orbitrap (60,000 resolution; maximum injection time 50 ms; AGC 4 × 10^5^). Charge states between 2 and 6 were required for MS2 analysis, and a 120 s dynamic exclusion window was used. Top 10 MS2 scans were performed in the ion trap with CID fragmentation (isolation window 0.5 Da; Rapid; NCE 35%; maximum injection time 50 ms; AGC 2 × 10^4^). An on-line real-time search algorithm (Orbiter) was used to trigger MS3 scans for quantification (*87*). MS3 scans were collected in the Orbitrap using a resolution of 50,000, NCE of 55%, maximum injection time of 200 ms, and AGC of 3.0 × 10^5^. The close out was set at two peptides per protein per fraction.

### Data analysis

Raw files were converted to mzXML, and monoisotopic peaks were re-assigned using Monocle (*88*). Searches were performed using the Comet search algorithm against a human database downloaded from Uniprot in May 2021. A 50 ppm precursor ion tolerance, 1.0005 fragment ion tolerance, and 0.4 fragment bin offset for MS2 scans were collected in the ion trap. TMT on lysine residues and peptide N-termini (+229.1629 Da) and carbamidomethylation of cysteine residues (+57.0215 Da) were set as static modifications, while oxidation of methionine residues (+15.9949 Da) was set as a variable modification. Each run was filtered separately to a 1% False Discovery Rate (FDR) on the peptide-spectrum match (PSM) level. Then proteins were filtered to the target 1% FDR level across the entire combined data set. For reporter ion quantification, a 0.003 Da window around the theoretical m/z of each reporter ion was scanned, and the most intense m/z was used. Reporter ion intensities were adjusted to correct for isotopic impurities of the different TMTpro reagents according to manufacturer specifications. Peptides were filtered to include only those with a summed signal-to-noise (SN) ≥ 110 across all TMT channels. The signal-to-noise (S/N) measurements of peptides assigned to each protein were summed (for a given protein). These values were normalized so that the sum of the signal for all proteins in each channel was equivalent thereby accounting for equal protein loading.

#### Data and statistical analyses

Data were analyzed off-line using PulseFit (HEKA), FlowJo, Microsoft Excel, MATLAB, and GraphPad Prism software. Statistical analyses were performed in GraphPad Prism and Microsoft Excel using built-in functions (GraphPad Prism) and the Solver function (Microsoft Excel). Statistically significant differences between means (set at *P* < 0.05) were evaluated using specified statistical tests. Data are presented as means ± SEM.

## Supporting information

Compiled supplemental data

## Acknowledgements

We thank Ming Chen for technical support. This work was supported by grants RO1-HL121253, RO1-HL142111, R01-NS126850 and P01-HL164319 from the NIH (to HMC); F31 DK118866 (to T.J.M); AHA predoctoral fellowship 20PRE35210815 (to ADB); and AHA postdoctoral fellowship POST1019343 (to SSK). Flow cytometry experiments were performed in the CCTI Flow Cytometry Core, supported in part by the NIH (S10RR027050). Confocal images were collected in the HICCC Confocal and Specialized Microscopy Shared Resource, supported by NIH (P30 CA013696).

## Author Contributions

TJM, ADB, EA, OBC, HMC designed experiments; TJM, ADB, EA, SSK, XZ, PC, MD performed experiments; TJM, ADB, EA, SSK, XZ, MD, OBC, HMC analyzed data; TJM, ADB, EA, OBC, HMC wrote the paper; TJM, ADB, EA, RSK, OBC, HMC edited the paper; TJM, ADB, RSK, HMC obtained funding.

