## Supplementary material for "Ion channel inhibition by targeted recruitment of NEDD4-2 with divalent nanobodies": Compiled supplemental data

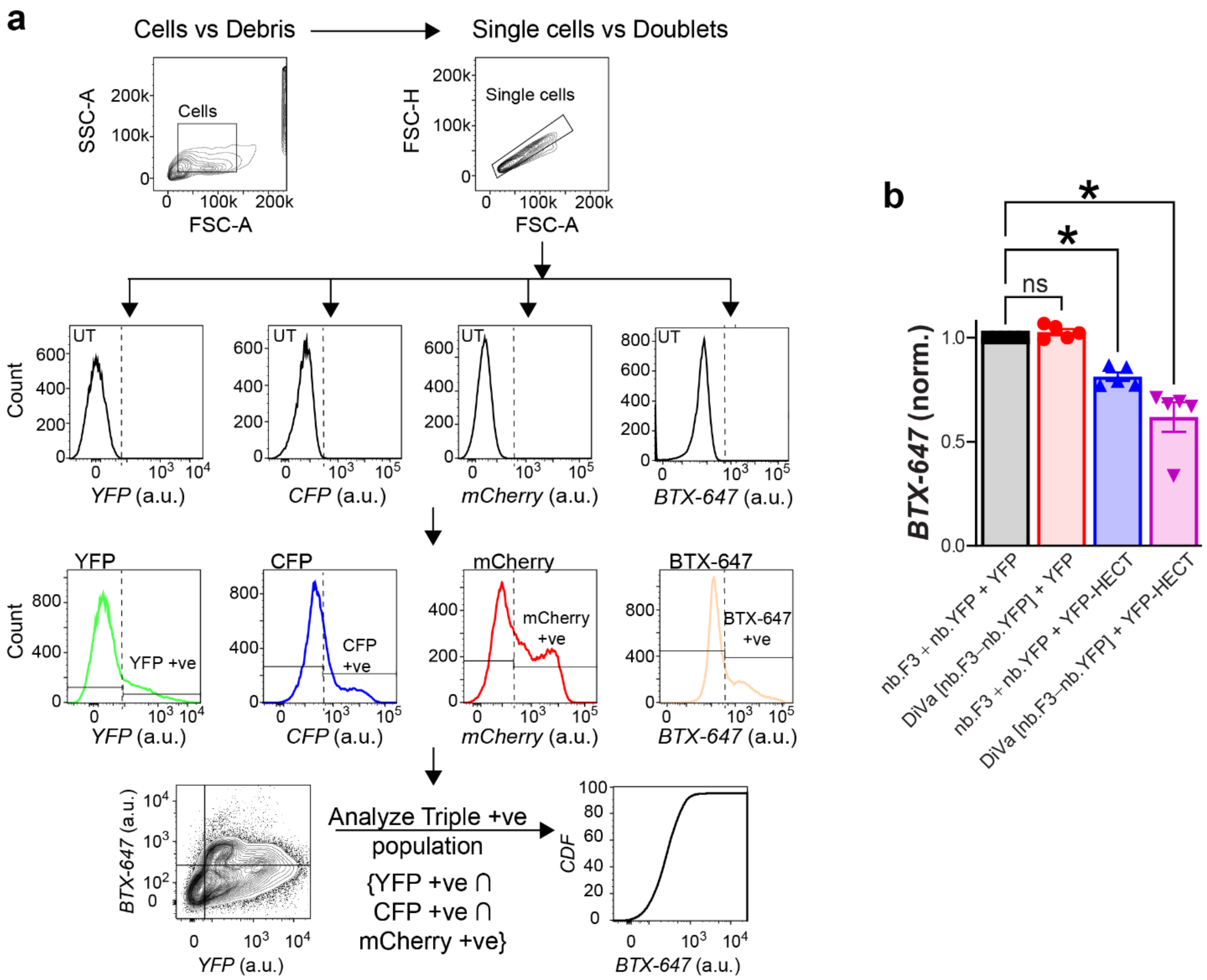


**Supplemental Figure 1. Targeted recruitment of NEDD4-2 HECT Domain with divalent nanobodies reduces α1B cell surface expression**

(a) Flow cytometry gating scheme used for experiments described in Figure 1. a.u. implies arbitrary units. (b) Aggregate data from flow cytometry cell-surface labeling experiments showing the effects of the indicated experimental conditions on $\alpha_1$B cell surface expression. For each experimental condition, data points are geometric means of the fluorescence intensities from ≥ 5000 cells normalized to the control group (black bar). Averages are means ± SEM. *$p<0.05$ compared to control by One-way ANOVA test with Tukey's multiple comparisons test; n.s implies not significantly different from control.

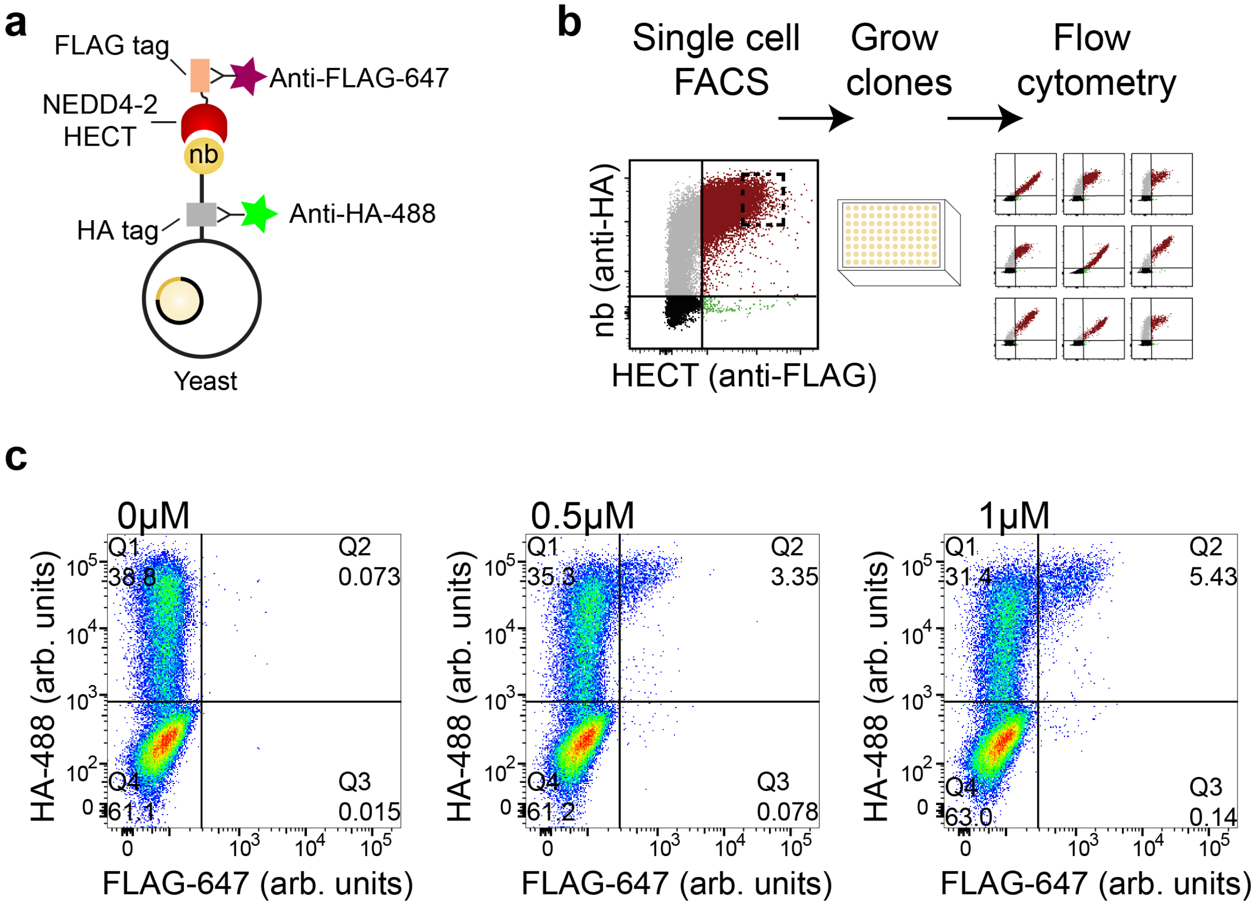


**Supplemental Figure 2. Identification of nanobody binders to NEDD4-2 HECT domain**
(a) Schematic showing the expression of HA and FLAG tags by the yeast nanobodies and NEDD4-2 HECT domain respectively. (b) Flow chart depicting the process leading to the isolation of NEDD4-2 HECT domain nanobody binders. (c) Exemplar flow cytometry dot plots showing enrichment in the population of nanobody binders (in quadrant 2; Q2) using 0μM, 0.5μM, or 1μM of purified NEDD4-2 HECT domain. HA-tagged nanobodies and FLAG-tagged NEDD4-2 HECT domain were detected by Alexa-fluor 488 anti-HA and Alexa-fluor 647 anti-FLAG antibodies respectively.

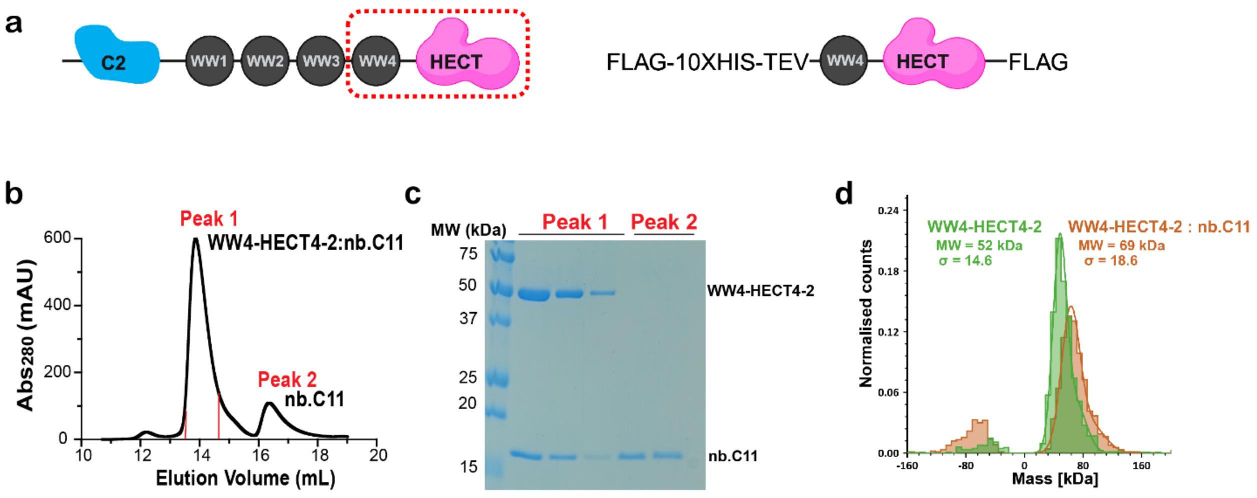


**Supplemental Figure 3. Purification and Biochemical characterization of WW4-HECT4-2:nb.C11 Complex**

(a) Left, Schematic showing the modular domains of NEDD4-2 E3 ligase. NEDD4-2 consist of N-terminal C2 domain (blue), four WW domains (black), and a HECT ubiquitin ligase domain (magenta). Shown in red dotted rectangle is the optimized construct we used for structural studies. The inclusion of proximal WW4 was to stabilize and add extra mass to the HECT domain. Right, Construct design used for expression and purification of WW4-HECT of NEDD4-2 from E.coli. (b) Gel filtration profile of WW4-HECT4-2:nb.C11 complex used for freezing grids. (c) Coomassie-stained SDS gel of peak 1 and peak 2. Fractions of peak 1 corresponding to WW4-HECT:nb.C11 complex in B were collected and concentrated for Cryo-EM studies. (d) Refeyn Mass Photometry Profiles of WW4-HECT4-2 and WW4-HECT4-2:nb.C11 complex (used for freezing on grids) showing a shift in molecular weight (MW) to the right indicating a larger MW complex (69 kDa) when compared to WW4-HECT4-2 alone (52 kDa).

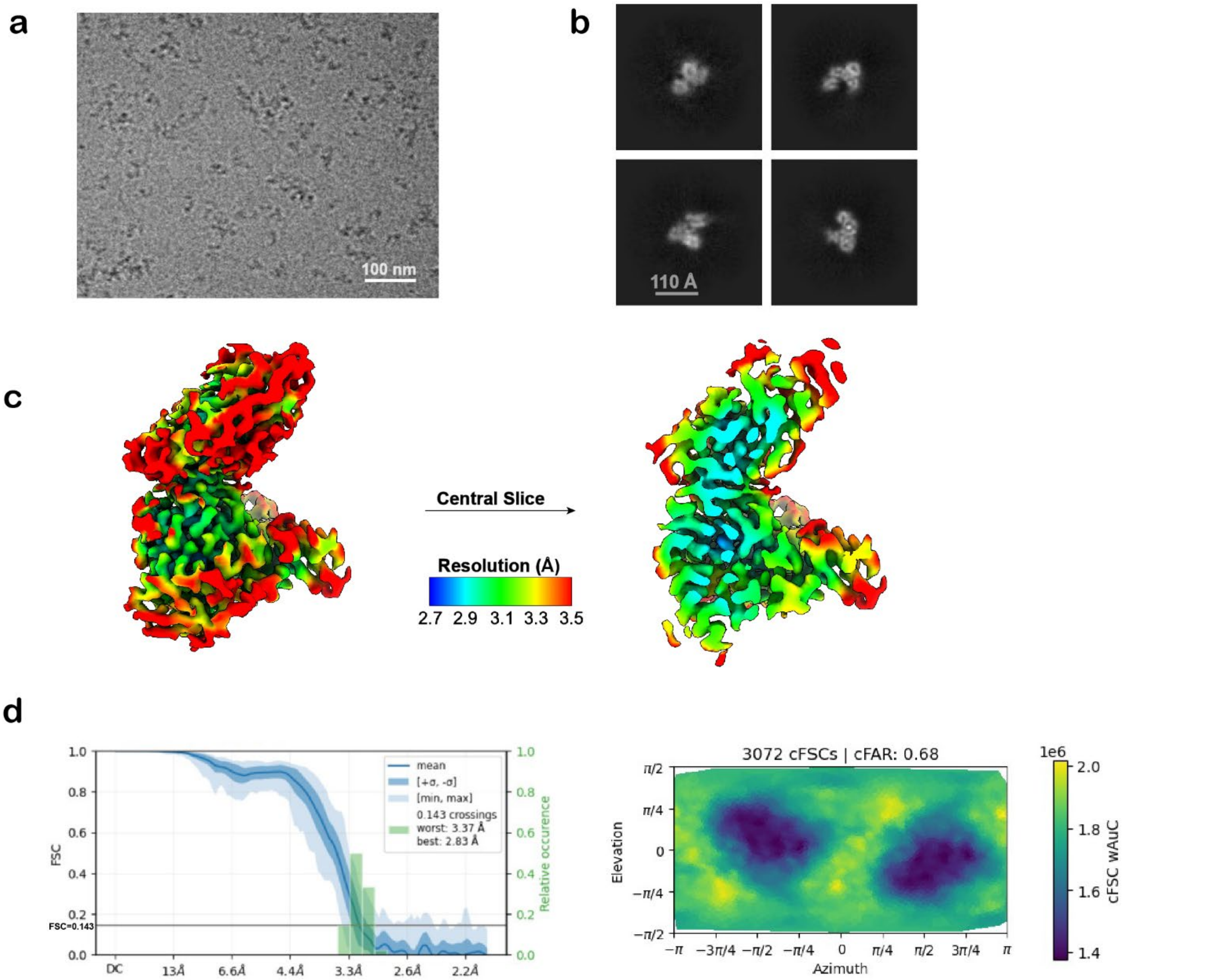


**Supplemental Figure 4. Cryo-EM Analysis of WW4-HECT4-2:nb.C11 Complex**

(a) Representative micrograph showing particle distribution for WW4-HECT4-2:nb.C11 complex. 7,790 micrographs were recorded in total (b) Representative 2D class averages showing high resolution features of the complex. (c) Two views of the final map colored according to local resolution (scale on the bottom). Local resolution was calculated in CryoSPARC 4.4. Sharpened density maps are contoured at 0.13. (d) Right, 3D FSC curves and preferred orientation analysis. The blue line indicates the global FSC. The FSC calculations used mask shown in Figure 7. Left, Orientation distribution analysis on the final refined map showing density isotropy and directional overall resolution across the map. The spatial distribution of particles in the final iteration of 3D refinement as calculated in CryoSPARC 4.4

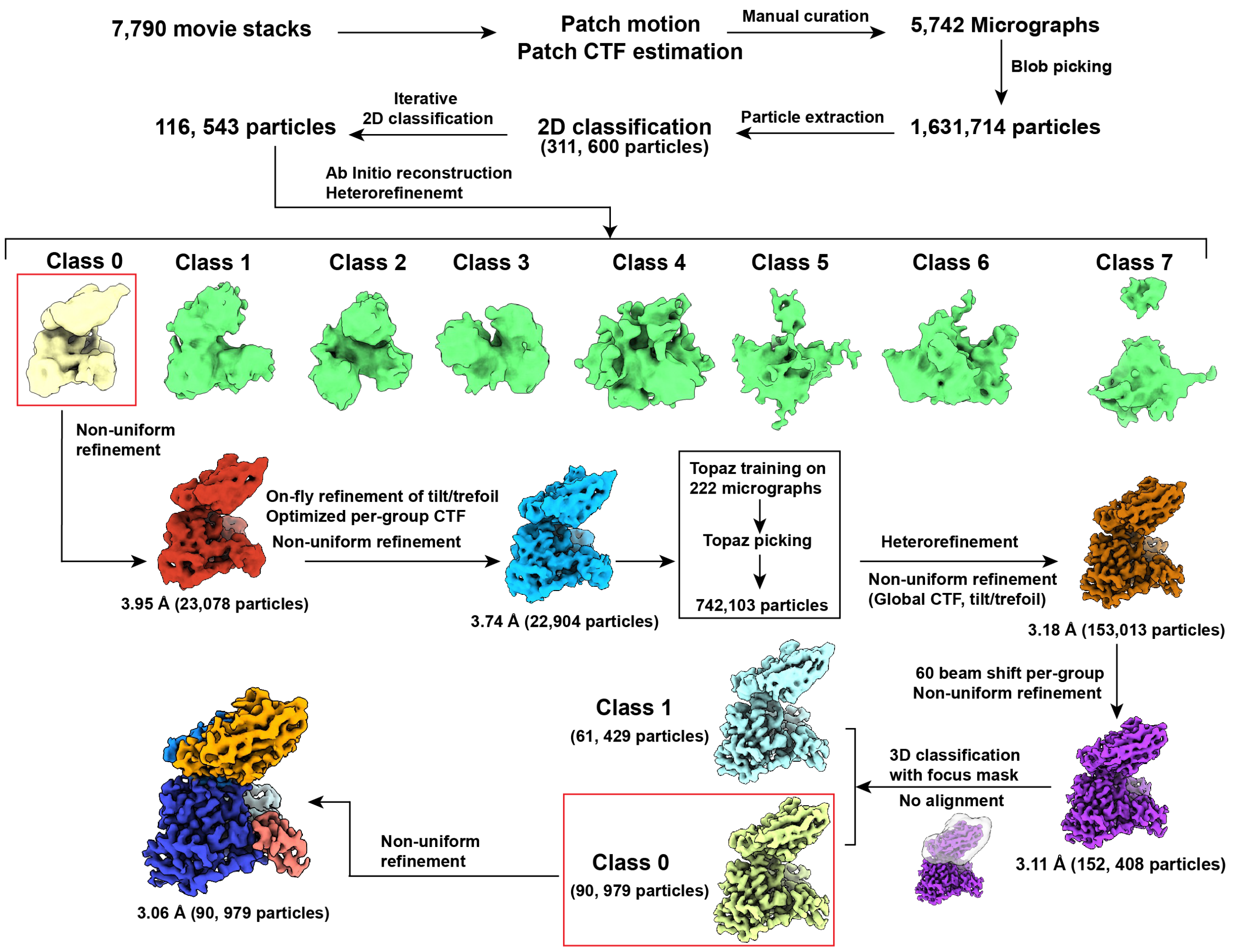


**Supplemental Figure 5. Flow chart for EM processing and refinement WW4-HECT:nb.C11 Complex.** All processing was performed in cryoSPARC 4.4. (See Methods for details).

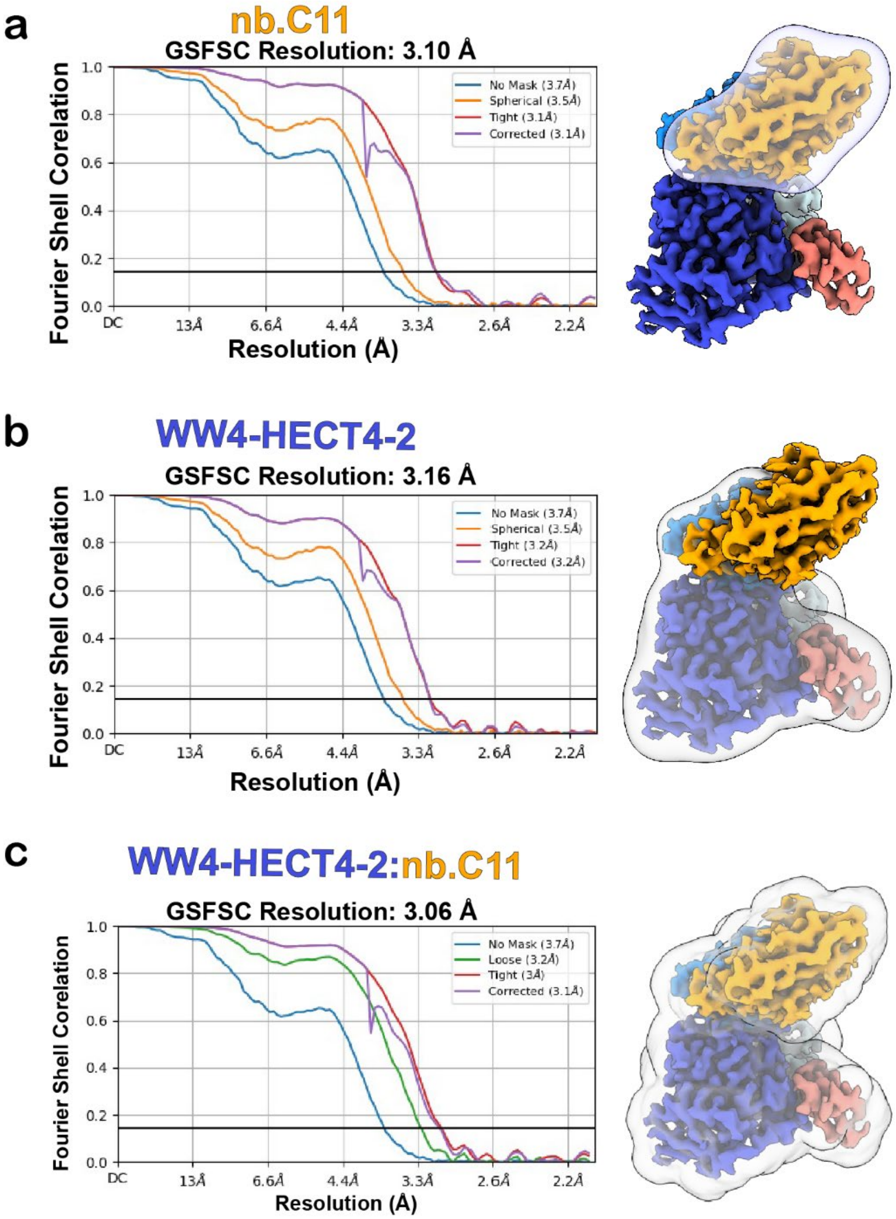


**Supplemental Figure 6. Data Quality and overall resolution for WW4-HECT4-2:nb.C11 Complex**
(a) Left, Fourier Shell Correlation (FSC) curve for overall resolution of the nb.C11. Right, A map of WW4-HECT4-2:nb.C11 complex showing the mask used for FSC calculation. (b). Left, Fourier Shell Correlation (FSC) curve for overall resolution of the WW4-HECT4-2. Right, A map of WW4-HECT4-2:nb.C11 complex showing the mask used for FSC calculation. (c) Left, Fourier Shell Correlation (FSC) curve for the overall resolution of the WW4-HECT4-2:nb.C11 Complex. Right, A map of WW4-HECT4-2:nb.C11 showing the mask used for FSC calculation.

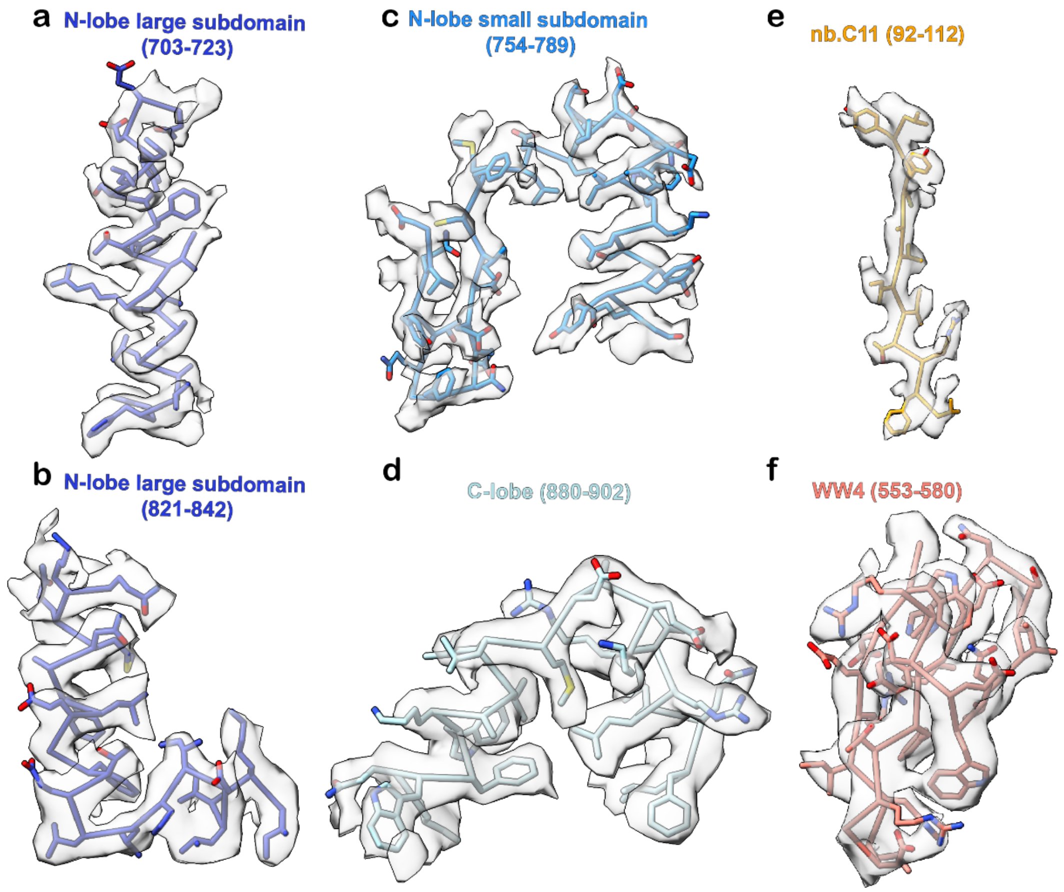


**Supplemental Figure 7. EM map quality assessment of WW4-HECT4-2:nb.C11 Complex**
Map and fitted model for different segments of the N-lobe subdomain (a,b, c), C-lobe of NEDD4-2 HECT domain (d), nb.C11 (e) and WW4 (f). Sharpened density maps are contoured at 0.136.

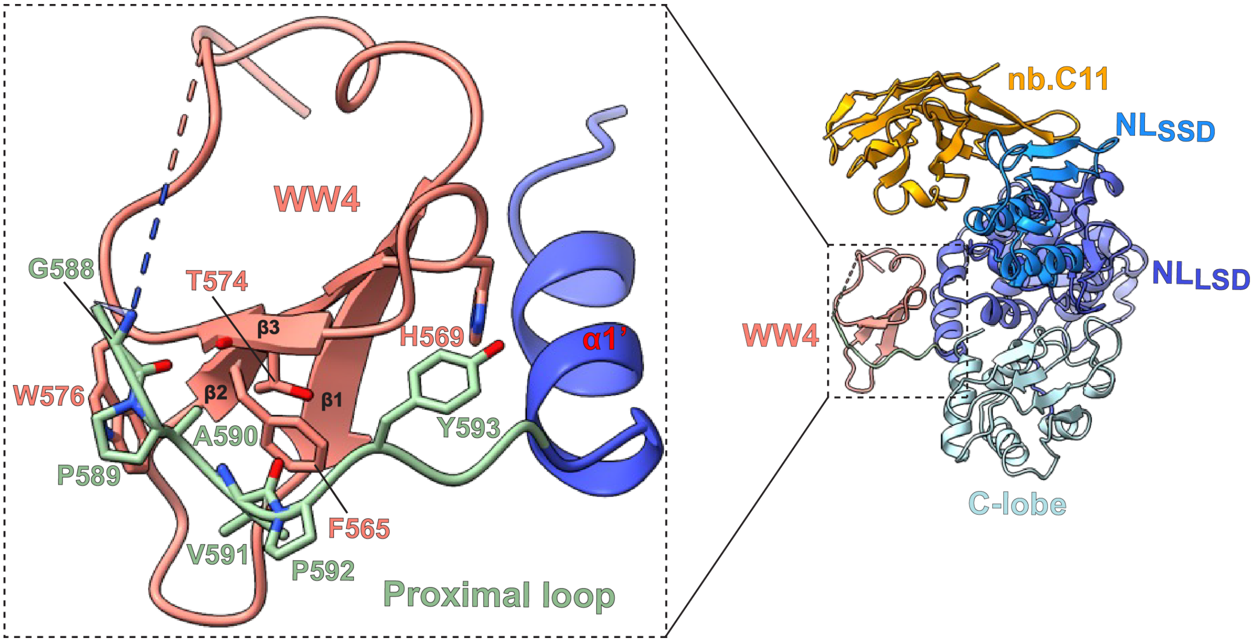


**Supplemental Figure 8. Close-up view of intramolecular interaction of WW4 with HECT4-2 proximal loop**

Structural analysis of the WW4:HECT4-2 interface including the close-up showing the interaction of the proximal loop (light green) and WW4. WW4 (Salmon) folds into three canonical anti-parallel β-strands to have an ordered interaction with the proximal loop which connects WW4 to the α1 helix at the start of HECT4-2 domain. The proximal loop utilizes P589, V591, P592, and Y593 for this stabilizing interaction, which constitute a non-canonical PAVPY (PY) motif not previously reported in literature for this class of enzyme.

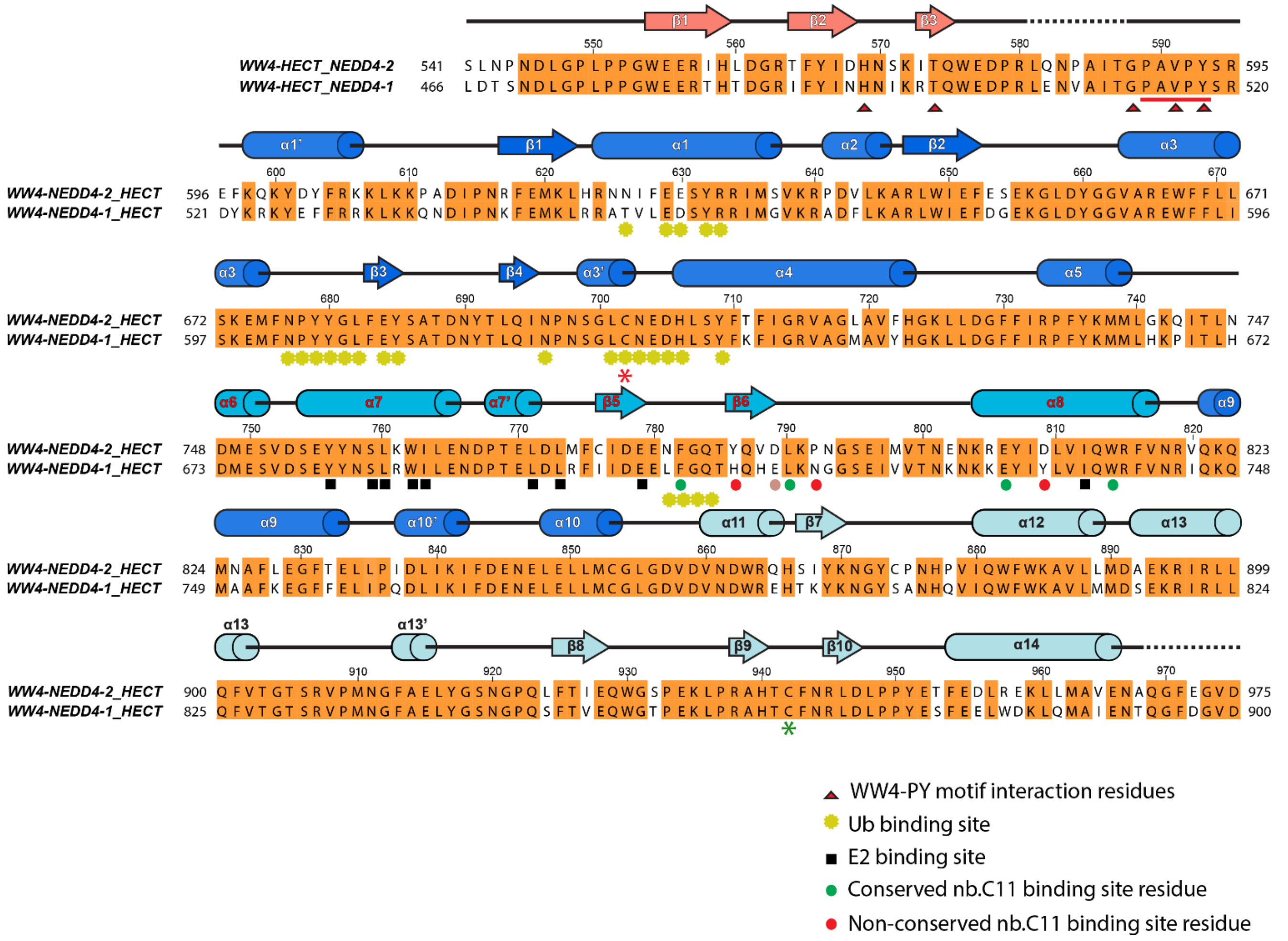


**Supplemental Figure 9: Sequence alignment and secondary structure of NEDD4-2 and NEDD4-1 HECT domains.** NEDD4-1 is close homolog of NEDD4-2 with 83 % sequence similarity between their HECT domains. The alignment was produced by Muscle 3.8 and manually curated in Jalview. Residues with absolute conservation are shaded in orange. Key residues of the HECT4-2 important for nb.C11 binding based on the structural information are indicated in dotted circles (green circles, conserved residues; red circles, non-conserved residues; brown circle, semi-conserved residue). Residues important for E2 binding are indicated in black squares, and those engaging in the WW4-PY motif interaction are represented by red triangles. The binding site for ubiquitin on the N-lobe is highlighted in yellow diamond. The strictly conserved catalytic cysteine is highlighted in green asterisk while non-catalytic cysteine is highlighted in red asterisk. Residues numbering corresponds to canonical human FL-NEDD4-2 protein (Uniprot ID: Q96PU5-1). The secondary structural elements are illustrated with α-helices and β-sheets as arrows. Dotted line indicate that residues were not visible in the electron density. Residue numbering correspond to canonical full-length human NEDD4-2 (Uniprot Id: Q96PU5-1) sequence.

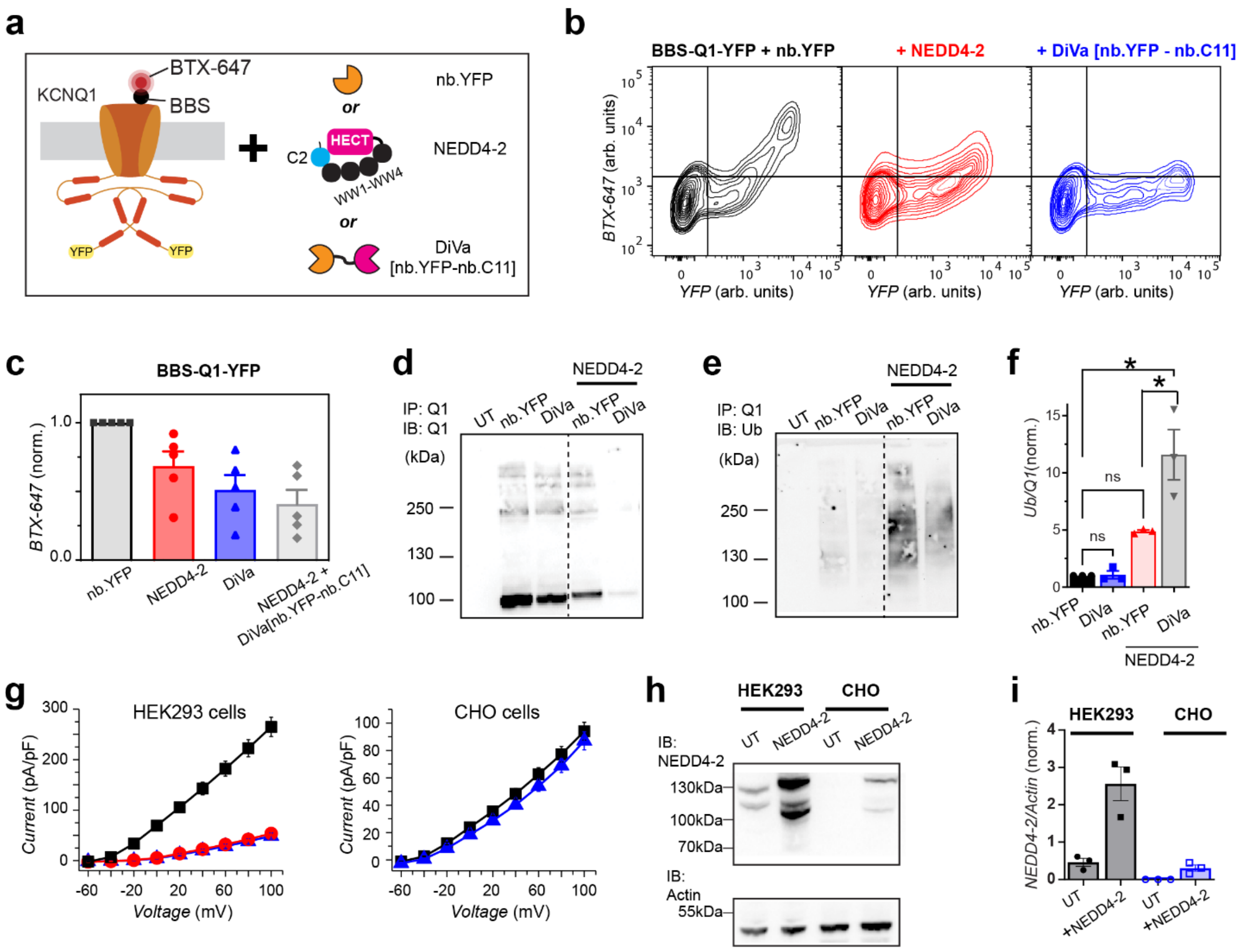


**Supplemental Figure 10. Mechanism of KCNQ1 inhibition by targeted NEDD4-2 recruitment**

(a) Cartoon of the YFP-tagged KCNQ1 channel expressing an extracellular epitope tag (bungarotoxin binding site, BBS). Alexa-fluor 647-conjugated bungarotoxin (BTX) binding to the BBS allows for the labeling of the cell-surface channel population in non-permeabilized transfected HEK293 cells during flow cytometry experiments. (b) Exemplar flow cytometry CDF plots showing the effects of over-expression of nb.YFP (black); NEDD4-2 (red), or DiVa [nb.YFP-nb.C11] (blue) on co-transfected BBS-KCNQ1-YFP in HEK293 cells. n≥5000 cells per condition. (c) Aggregate data showing the effects of over-expression of the indicated DNA constructs on co-transfected BBS-KCNQ1-YFP in HEK293 cells. For each experimental condition, data points are geometric means of the fluorescence intensities from ≥ 5000 cells normalized to the control group (black bar). Averages are means ± SEM. (d-e) Exemplar immunoblots of immunoprecipitated KCNQ1-YFP showing the effects of the indicated conditions on KCNQ1 steady-state ubiquitination. (f) Aggregate data quantification of changes in KCNQ1 ubiquitination. Ubiquitin expression for each condition was divided by the corresponding Q1 expression for that condition. The resulting numbers for each of the indicated treatment groups were normalized to that of the BBS-KCNQ1-YFP + nb.YFP control group (black). *$p<0.05$ compared to nb.F3 control by One-way ANOVA test with Tukey's multiple comparisons test. (g) Population I-V curves evoked from HEK293 (left; reproduced from Fig. 4c) and CHO (right) cells co-transfected with KCNQ1-YFP and nb.YFP (black); NEDD4-2 (red) or DiVa [nb.YFP-nb.C11] (blue). For experiments in CHO cells, n=10 cells per condition. (h) Representative immunoblot (h) and quantification (i) showing NEDD4-2 expression in transfected and untransfected (UT) HEK293 and CHO cells.

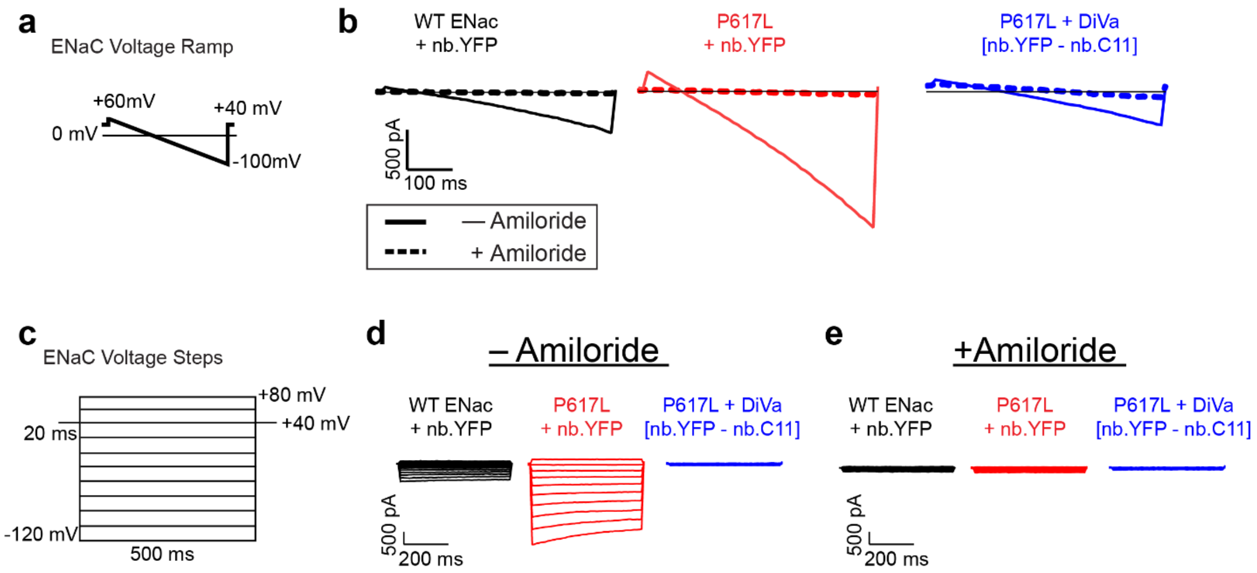


**Supplemental Figure 11. Inhibition of amiloride-sensitive ENaC currents by targeted NEDD4-2 recruitment**

a) Schematic of the voltage-ramp protocol used for ENaC whole-cell current measurements.

b) Exemplar voltage-ramp current traces showing ENaC currents in the presence (dashed lines) and absence (solid lines) of 10μM amiloride for the indicated experimental conditions. Each trace is a representative trace at steady-state for the specified condition.

c) Schematic of the voltage-step protocol used for ENaC whole-cell current measurements.

d-e) Exemplar whole-cell current traces evoked from HEK293 cells transfected as indicated in the absence (d) and presence (e) of 10μM amiloride.

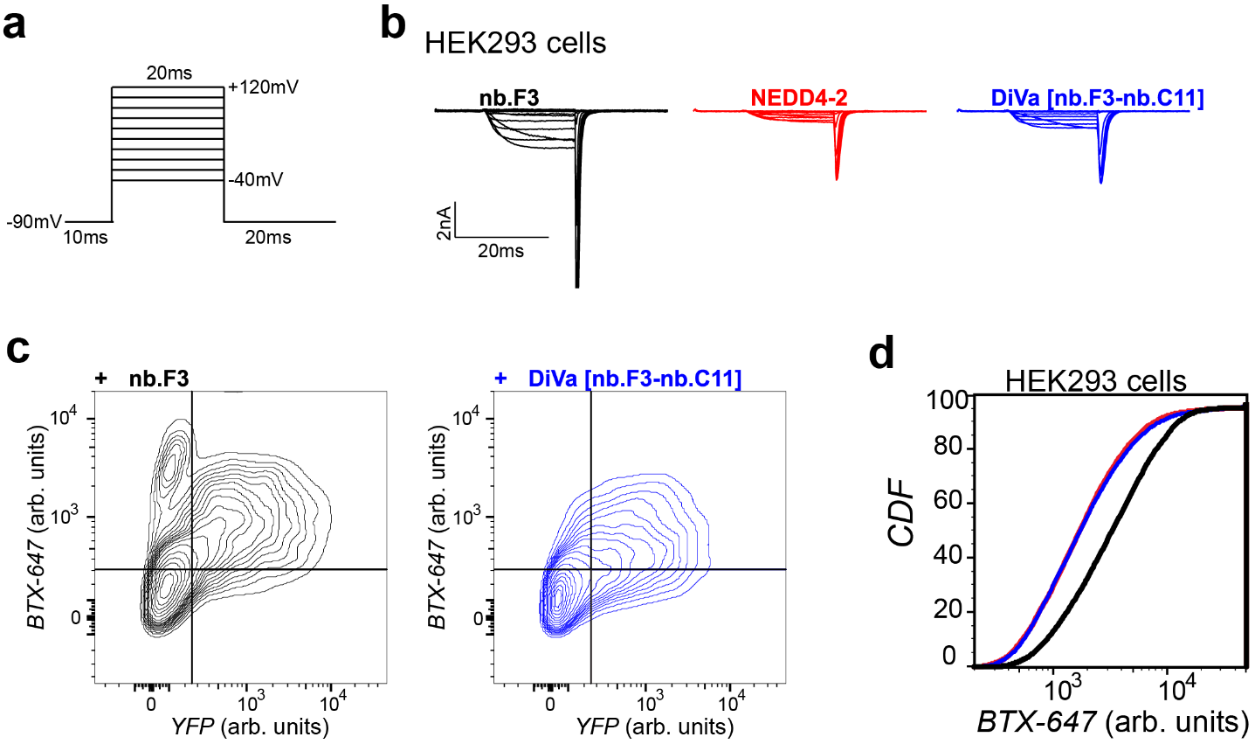


**Supplemental Figure 12. Inhibition of CaV2.2 by targeted NEDD4-2 recruitment**

(a) Schematic of the voltage-step protocol used for $Ca_V2.2$ whole-cell current measurements. (b) Exemplar whole-cell current traces evoked from HEK293 cells transfected with $\alpha_1$B, YFP-β2a, α2δ1-P2a-mCherry and nb.F3 (black); NEDD4-2 (red); or DiVa[nb.F3-nb.C11]. (c) Exemplar flow cytometry contour plots showing the effects of the indicated conditions on the surface expression of $\alpha_1$B $Ca_V2.2$ subunit in transiently transfected HEK293 cells. (d) Exemplar CDF plots showing the effects of co-transfection of DiVa[nb.F3-nb.C11] (blue); NEDD4-2; or DiVa[nb.F3-nb.C11] + NEDD4-2 (grey) on BBS-KCNQ1-YFP relative to the control group (BBS-KCNQ1-YFP + nb.YFP; black). a.u., arbitrary units

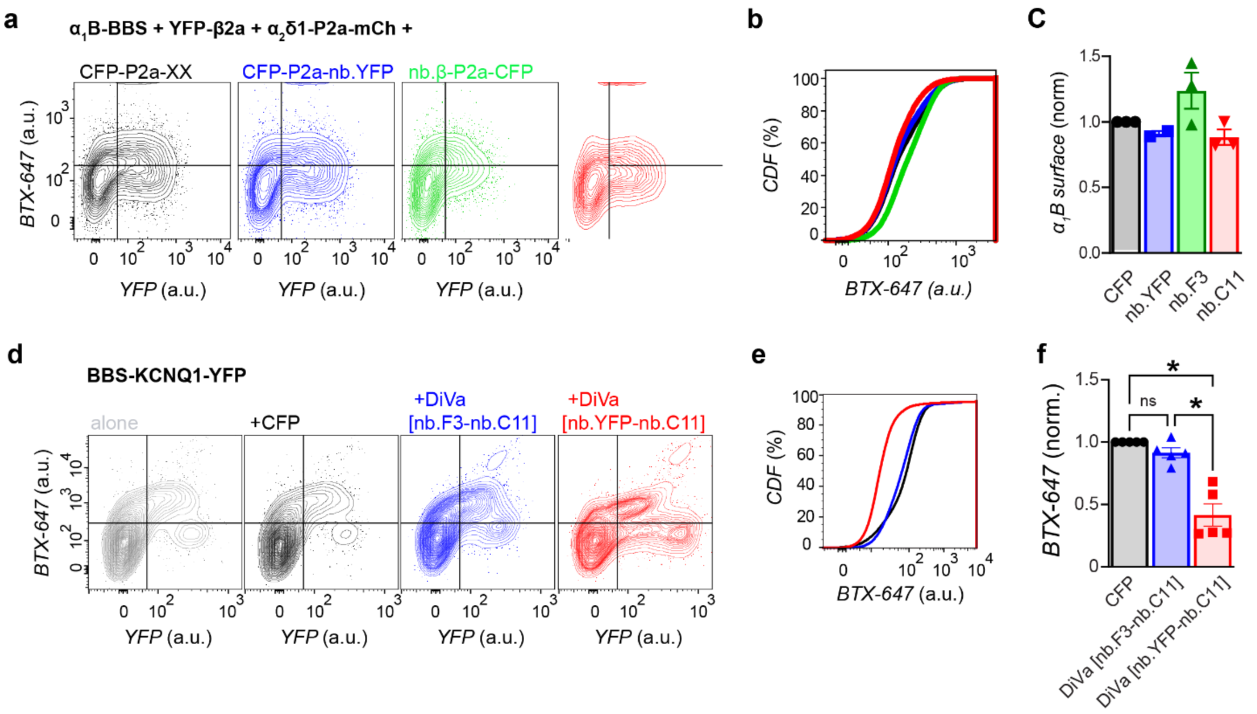


**Supplemental Figure 13. DiVa[nb.F3-nb.C11] selectively inhibits $Ca_V2.2$ channel complex, but not KCNQ1.**

(a-b) Exemplar flow cytometry contour (a) and CDF (b) plots showing the effects of the indicated conditions on the surface expression of the $\alpha_1$B subunit of the $Ca_V2.2$ channel complex. The traces represent HEK293 cells transfected with $\alpha_1$B-BBS, YFP-β2a, α2δ1-P2a-mCherry and CFP-P2a (black); CFP-P2a-nb.YFP (blue); nb.F3-P2a-CFP (green); or CFP-P2a-nb.C11 (red).

(c) Aggregate data from flow cytometry cell-surface labeling experiments showing the effects of the indicated experimental conditions on $\alpha_1$B cell surface expression. For each experimental condition, data points are geometric means of the fluorescence intensities from ≥ 5000 cells normalized to the control group (black bar). Averages are means ± SEM. (d-e) Exemplar flow cytometry contour (d) and CDF (e) plots showing the effects of the indicated conditions on the surface expression of the KCNQ1 channel. In (e), the traces represent HEK293 cells transfected with BBS-KCNQ1-YFP and CFP-P2a (black); CFP-P2a-DiVa[nb.F3-nb.C11] (blue); or CFP-P2a-DiVa [nb.YFP-nb.C11] (red). (f) Aggregate data from flow cytometry cell-surface labeling experiments showing the effects of the indicated experimental conditions on KCNQ1 cell surface expression. For each experimental condition, data points are geometric means of the fluorescence intensities from ≥ 5000 cells normalized to the control group (CFP; black bar). Averages are means ± SEM. *$p<0.05$ compared to CFP control by One-way ANOVA test with Tukey's multiple comparisons test.

**Supplementary Table S1: Cryo-EM data collection, refinement and validation statistics**

| | **WW4-HECT4-2:nb.C11 Complex**<br>PDB: XXXX<br>EMDB: XXXX |
|---|---|
| **Data Collection** | |
| Microscope | Titan Krios |
| Detector | Gatan K3 (Counting) |
| Energy Filter, slit | 20 eV |
| Magnification | 165,000x |
| Volage (kV) | 300 |
| Electron Exposure ($e^-/Å^2$) | 109.2 |
| Exposure time (s) | 1.8 |
| Dose Rate ($e^-$/pixel/s) | 16.1 |
| Frame Number | 100 |
| Defocus range (µm) | -2.5 to -1 |
| Pixel size (Å) | 0.5135 |
| Number of micrographs collected | 7,790 |
| **3D Reconstruction** | |
| Micrographs used | 5,742 |
| Symmetry imposed | C1 |
| Initial particle images (no.) | 1,631,714 |
| Final particle images (no.) | 90,979 |
| Map resolution (Å) | 3.06 |
| FSC threshold | 0.143 |
| Local resolution range | 2.7-3.5 |
| **Model Refinement** | |
| Initial Model (PDB Code) | 4BE8, 2MPT |
| Model composition | |
| Non-hydrogen atoms | 42880 |
| Protein residues | 539 |
| Ligands | 0 |
| R.m.s deviations | |
| Bond length (Å) | 0.003 |
| Bond Angles (°) | 0.538 |
| Validation | |
| Molprobity score | 1.95 |
| Clash score | 5.85 |
| Poor rotamers (%) | 0 |
| Ramachandra Statistics (%) | |
| Outliners | 0.00 |
| Allowed | 2.63 |
| Favored | 97.37 |
